# BANKSY: A Spatial Omics Algorithm that Unifies Cell Type Clustering and Tissue Domain Segmentation

**DOI:** 10.1101/2022.04.14.488259

**Authors:** Vipul Singhal, Nigel Chou, Joseph Lee, Jinyue Liu, Wan Kee Chock, Li Lin, Yun-Ching Chang, Erica Teo, Hwee Kuan Lee, Kok Hao Chen, Shyam Prabhakar

## Abstract

Each cell type in a solid tissue has a characteristic transcriptome and spatial arrangement, both of which are observable using modern spatial omics assays. However, the common practice is still to ignore spatial information when clustering cells to identify cell types. In fact, spatial location is typically considered only when solving the related, but distinct, problem of demarcating tissue domains (which could include multiple cell types). We present BANKSY, an algorithm that unifies cell type clustering and domain segmentation by constructing a product space of cell and neighbourhood transcriptomes, representing cell state and microenvironment, respectively. BANKSY’s spatial kernel-based feature augmentation strategy improves performance and scalability on both tasks when tested on FISH-based and sequencing-based spatial omics data. Uniquely, BANKSY identified hitherto undetected niche-dependent cell states in two mouse brain regions. Lastly, we show that quality control of spatial omics data can be formulated as a domain identification problem and solved using BANKSY. BANKSY represents a biologically motivated, scalable, and versatile framework for analyzing spatial omics data.

## 1 Introduction

A fundamental biological property of solid tissues is the arrangement of individual cell types in stereotypical spatial patterns. With the emergence of spatial omics technologies, we can now study tissue structure by examining both the spatial locations and molecular profiles of cells. These technologies provide highly multiplexed transcriptomic, genomic or proteomic profiles of single cells, together with their spatial coordinates (for example: multiplexed FISH [1, 2], Slide-seq [3], Slide-DNA-seq [4], MIBI-TOF [5]), and promise to provide unprecedented insights into cellular states, functions and interactions within the tissue context. As in the case of single cell RNA-seq (scRNA-seq), one of the primary spatial omics data analysis tasks is to cluster cells by similarity, with each cluster defining a distinct cell type or subtype. The most biologically relevant approach in this case would be to cluster cells using their transcriptomes as well as their spatial relationships. Surprisingly however, virtually all previous spatial omics studies have used clustering algorithms originally designed for single cell RNA-seq data, which ignore spatial information [2, 6, 7]. It is thus important to develop a biologically motivated formalism for incorporating spatial location in cell type clustering.

Recently, two algorithms have been proposed for incorporating spatial information into cell type clustering. The first [8] assumes that physically distant cells are less similar to each other. A limitation of this approach is that cells of the same type are often far apart, for example in the case of elongated structures such as epithelial layers and blood vessels, intercalated immune cells resident in tissues and intermingled neuronal and macroglial cells in the brain. In addition, cells of the same type are often spatially separated within repeating structures such as the two cerebral hemispheres or neuroepithelial buds within a brain organoid. The other method [9] adapts a Markov random field (MRF) framework for cell type annotation, combining each cell’s gene expression signature with the cluster membership identities of its physical neighbours. While this method attempts to model both the cell’s own transcriptome and its microenvironment composition simultaneously, the tool is not scalable to large spatial datasets, the underlying probabilistic generative model is complex, and it requires prior knowledge of the number of clusters.

In contrast to cell type clustering, multiple spatially informed algorithms have been developed to identify coherent domains in tissues (for example, cortical layers in mammalian brain). This is a distinct algorithmic problem, since each tissue domain could in principle include multiple distinct cell types. The earliest domain segmentation methods incorporated spatial information by encouraging physically proximal cells to have the same label (using Markov random fields) [10, 11]. One fundamental drawback of these methods is their assumption that an individual cell’s transcriptome resembles the average transcriptome of cells in its tissue domain. This assumption is not valid, since diverse cell types frequently reside within a single tissue domain. The other major family of tissue domain segmentation methods uses graph convolutional neural networks [12, 13], which are vulnerable to technical variability when the test dataset does not resemble training data.

Here, we introduce a biologically motivated strategy for combining molecular profiles with spatial locations, which addresses the above limitations. Our algorithm, named Building Aggregates with a Neighbourhood Kernel and Spatial Yardstick (BANKSY), leverages the fact that a cell within a tissue can be more fully represented by considering both its own transcriptome and the transcriptome of its local microenvironment. BANKSY uses a spatial kernel to compute the microenvironmental transcriptome as a weighted average over neighbouring cells. A major advantage of this strategy in the context of cell clustering is that it avoids the pitfall of assuming that cells of the same type or subtype are physically proximal. Another important advantage is that BANKSY can solve the distinct problems of cell type clustering and tissue domain segmentation within a unified feature augmentation framework.

The above strategy allows BANKSY to label spatially structured cell types and subtypes with high accuracy. In particular, BANKSY is adept at distinguishing subtly different cell subtypes residing in distinct microenvironments. Moreover, by modifying a single hyperparameter, BANKSY can be used to accurately detect tissue domains. Importantly, BANKSY’s strategy of feature augmentation by constructing a product space of own and neighbourhood transcriptomes allows it to leverage existing, highly scalable graph clustering solutions that can accommodate millions of cells [14, 15]. Finally, BANKSY is seamlessly inter-operable with the widely used bioinformatics pipelines Seurat (R), SingleCellExperiment (R), and Scanpy (Python). We anticipate that BANKSY’s biologically inspired approach for combining spatial and molecular data will yield rich insights into tissue architecture. Moreover, this approach can be further extended, potentially forming the basis for additional algorithms for analyzing spatial omics data.

## 2 Results

A direct approach to incorporating spatial information into clustering is to simply append cells’ spatial coordinates to the set of features defining their omic profiles (as was shown in Fig. 7 of [7]; see Supp. Section 1 for details). However, this approach is conceptually problematic because it causes cells in disparate physical locations to be labelled differently (Supp. Fig. 1), even when they have similar transcriptomic signatures, and should be assigned the same labels. Instead, a spatial clustering algorithm should not require that each distinct cell type be restricted to a single contiguous region, or even be near other cells of the same type. This situation can occur, for instance, when there are similar repetitive biological architectures present in a tissue or organ, as in the case of two cerebral hemispheres of a mouse brain (Fig. 3a, Supp. Fig 1), multiple neuroepithelial buds within a brain organoid (Fig. 5) or the intestinal crypts along the epithelium lining the colon.

**Figure 1.**
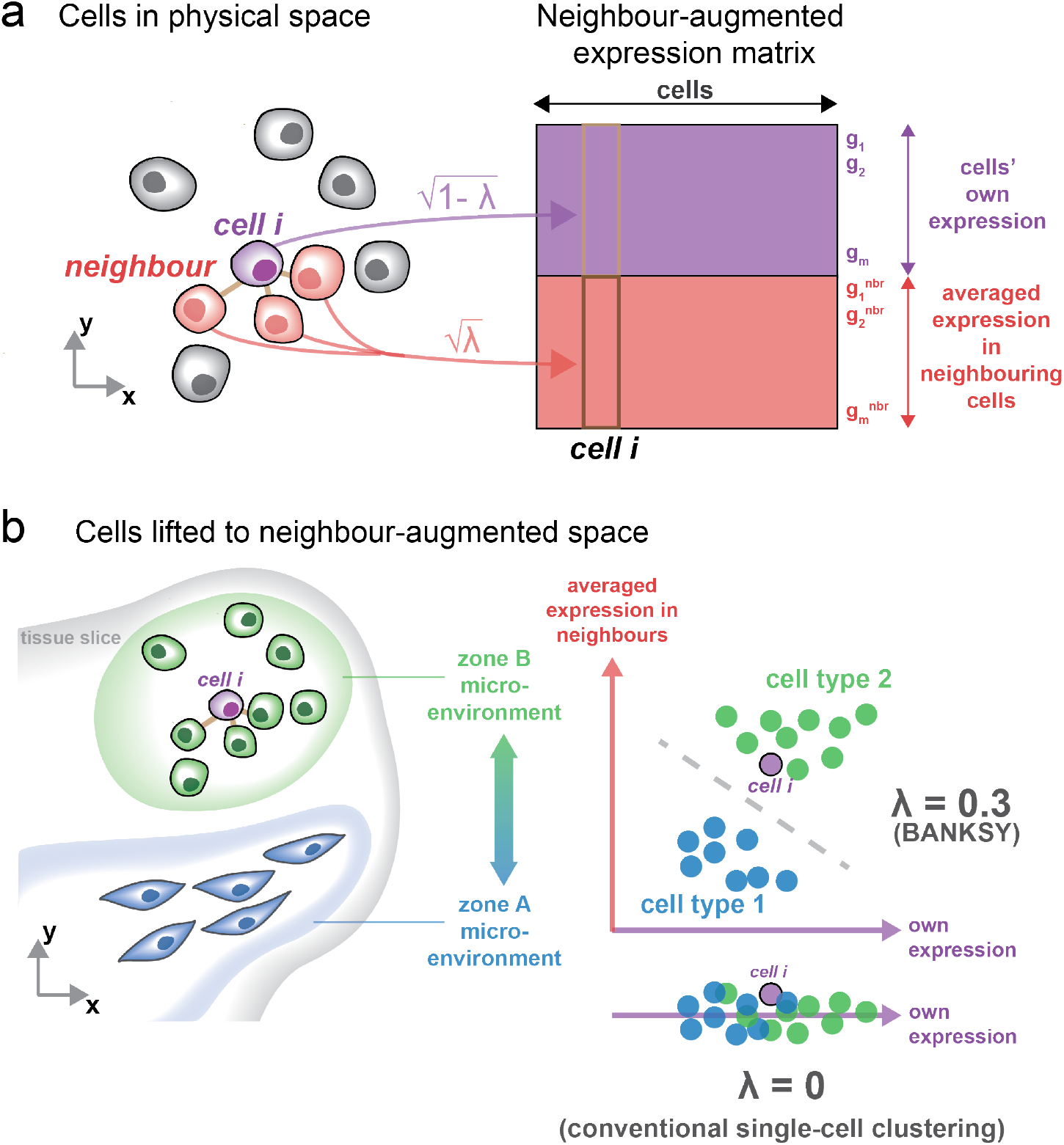
Schematic describing the incorporation of spatial microenvironment information into cell clustering. (a) The gene-cell matrix (purple) is augmented with a neighbourhood expression matrix (pink), where the column corresponding to the *i*^th^ cell is the averaged expression of that cell’s neighbours in physical space. (b) Visualisation of two cell types in the neighbour-augmented space. The averaged expression of neighbour cells, representing local microenvironment, helps to separate the two clusters which would be difficult to separate based on the cells’ own expression alone. For simplicity, we show microenvironments consisting of a single cell type (zone A and B containing cell types 1 and 2 respectively), but BANKSY is equally applicable to heterogeneous microenvironments with mixtures of cell types. For more details, see Supp. Fig. 2.

**Figure 2.**
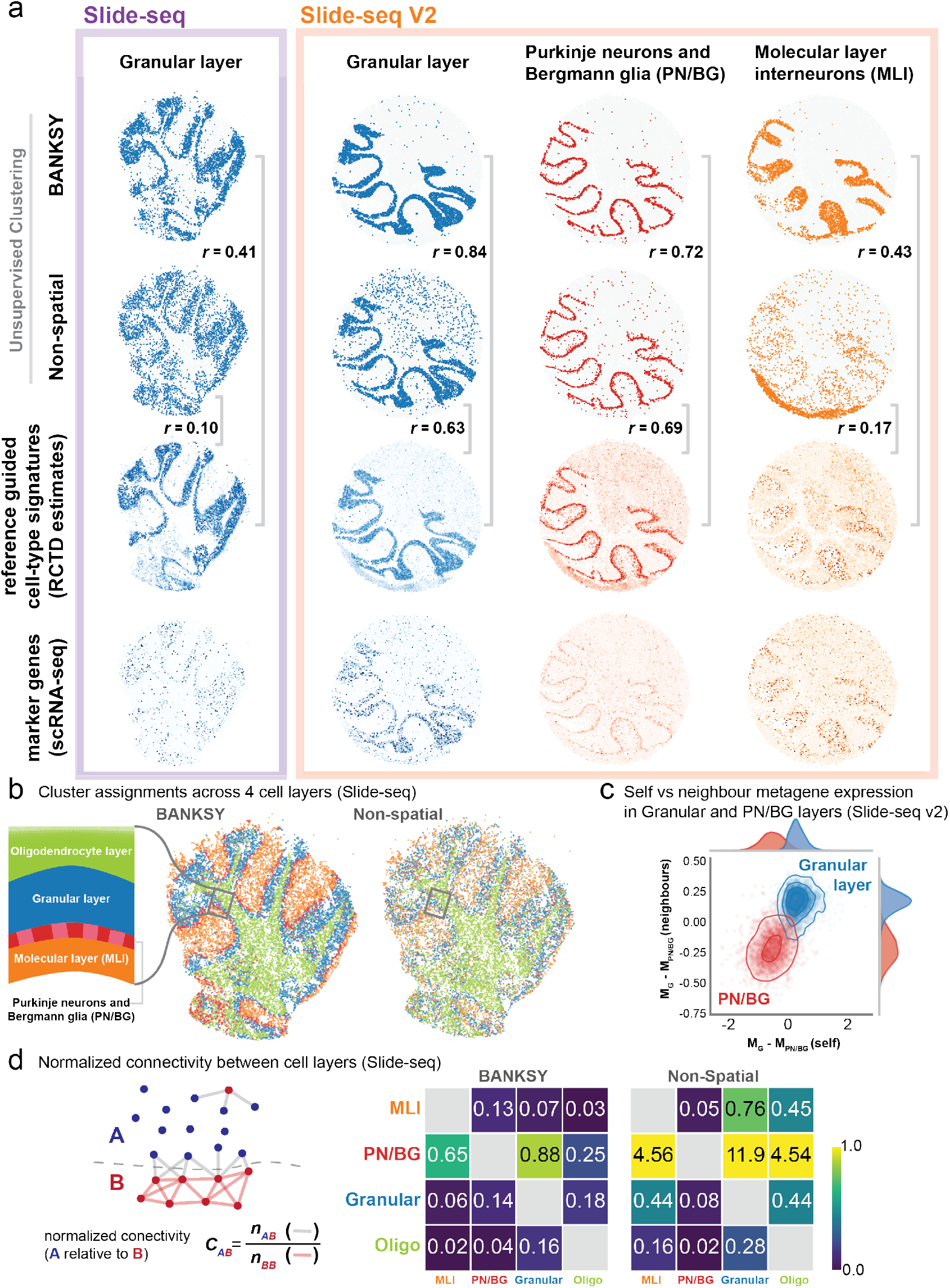
BANKSY refines cell type assignments in Slide-seq and Slide-seq v2 mouse cerebellum.: (a) Spatial map of three clusters corresponding to the granular layer (blue), the combined Purkinje neuron and Bergmann glia layer (PN/BG) (red) and molecular layer interneurons (MLI) (green) showing known layers of the mouse cerebellum. BANKSY clustering is compared to clustering without spatial information. Both are compared to robust cell type decomposition (RCTD, [3]) weights of the corresponding layer, with r values indicating the point-biserial correlation with RCTD weights. The bottom row shows the averaged expression of top differentially expressed (DE) marker genes from scRNA-Seq reference data. (b) Cell type assignments for four clusters in the Slide-seq mouse cerebellum (granular layer, PN/BG, MLI and the oligodendrocyte/polydendrocyte cluster). Schematic is adapted from [19] (c) Illustration of how neighbour gene expression (y-axis) enables cleaner separation of clusters. Each axis shows the difference in averaged expression of the top 20 DE genes (M_cluster_, Supp. Section 3) from the PN/BG cluster relative to the granular layer cluster (Slide-seq v2). Cells from granular layer cluster shown in blue, cells from PN/BG cluster in red. (d) Left: simplified schematic showing how normalised connection score between layers (A and B) was computed. Grey lines show edges between cells of different type, red lines show edges between cells of the same type. Right: normalised connection score matrix between layers in the Slide-seq dataset. Colourmap is clipped at 1. Higher numbers indicate greater intermingling between layers. Abbreviations: MLI– molecular layer interneurons, PN/BG–purkinje neurons and Bergmann glia, Granular–granular layer, Oligo–oligodendrocytes/polydendrocytes

**Figure 3.**
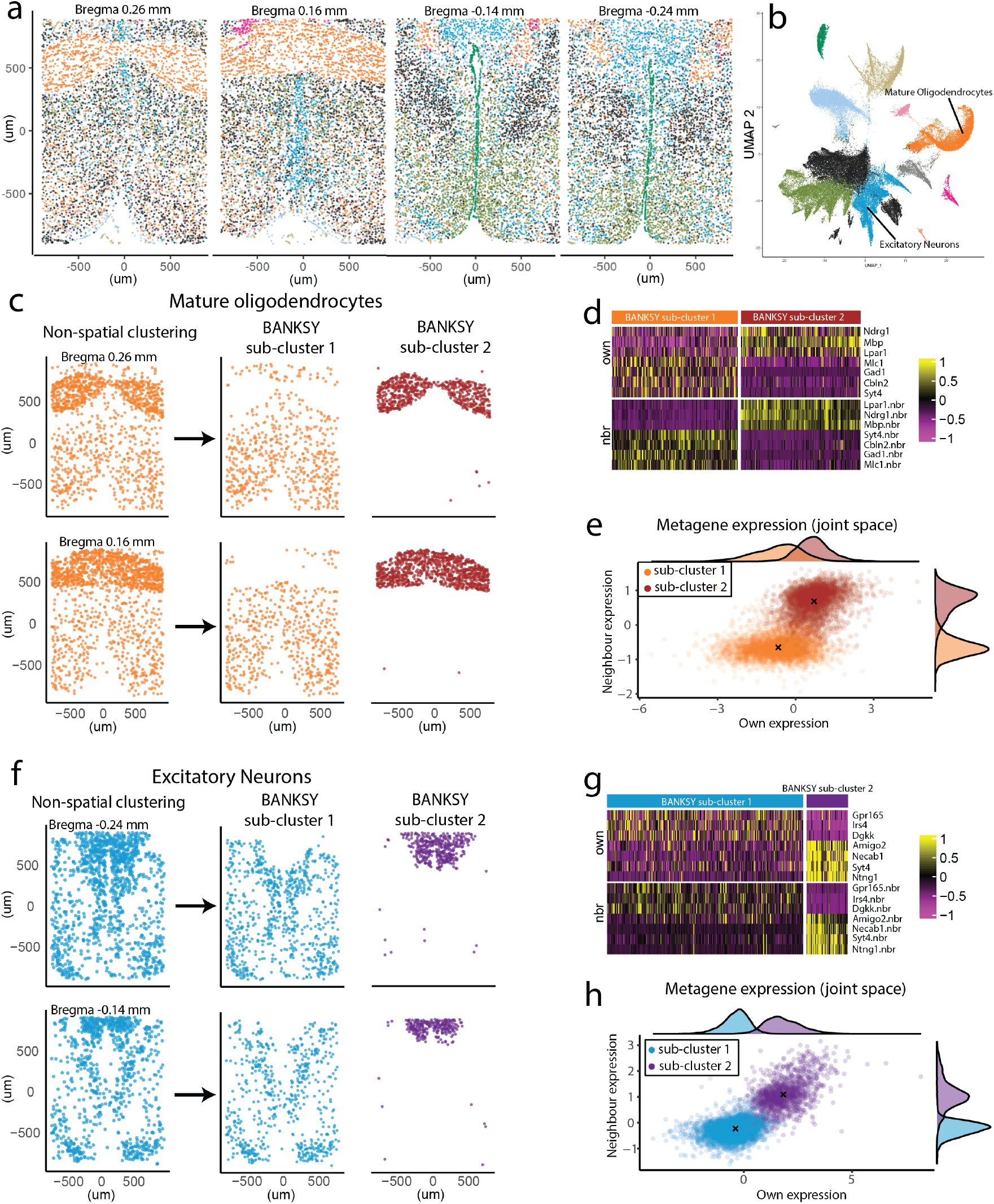
BANKSY finds oligodendrocyte and excitatory neuron cell subtypes in the mouse hypothalamus: (a) Overview of the mouse hypothalamus data from [2], coloured by non-spatial clustering labels (spatial tissue maps of 4 of the 12 z-slices in animal 1). (b) Corresponding UMAP computed using the gene expression values, and coloured by nonspatial (conventional) clustering colours. The mature oligodendrocytes and excitatory neurons are single clusters in non-spatial clustering. (c) Spatial maps for non-spatial vs BANKSY clustering show the physical locations of the two mature oligodendrocyte subclusters, two z-slices shown (Bregmas 0.26 mm and 0.16 mm). (d) Heatmap showing z-scaled expression values of DE genes distinguishing the two mature oligodendrocyte subclusters found by BANKSY. (e) Metagene constructed with the markers shown in (d), as described in Supp. Section 3. The same UMAP as (b), but coloured by the BANKSY colouring, can be found in Supp. Fig. 9a, and shows that the mature oligodendrocyte subclusters are intermixed in the own-expression space. (f-h) Analogous results for the excitatory neuron cluster.

**Figure 4.**
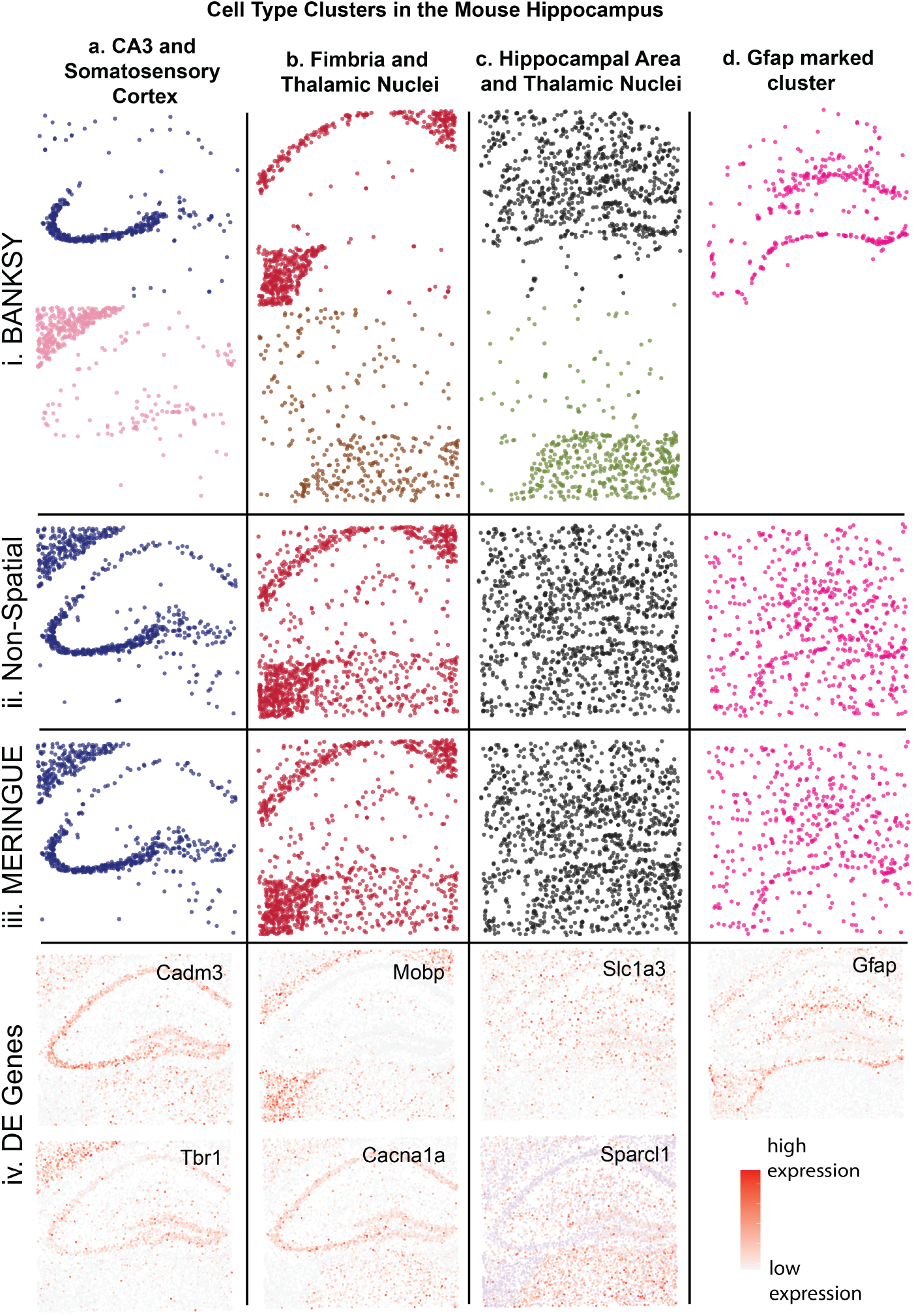
BANKSY applied to the mouse hippocampus reproduces effects similar to those found in Fig. 2 and 3. It leverages neighbourhood signature to distinguish clusters not separable by non-spatial clustering or MERINGUE [8], and is able to identify clusters more cleanly. Columns (a-d): Distinct clusters. Rows (i-iv): Distinct clustering methods and selected DE genes. (a, i) CA3 pyramidal neurons (blue) and cells in the somatosensory cortex (pink) were separated by BANKSY. (ii-iii) These clusters could not be resolved by non-spatial clustering and MERINGUE. Bottom blue box in Supp. Fig. 16 shows the neighbourhood expression of genes that distinguish these clusters, and Supp. Fig. 13a shows the genes that were differentially expressed between the own expressions of the cells in these two subclusters. (iv) Spatial distribution of representative DE genes (red: high expression, pale purple/white: low expression). (b, c) Two more pairs of clusters separated by BANKSY, but not by nonspatial clustering or MERINGUE. (i) Cluster assignment for BANKSY, (ii) non-spatial clustering, (iii) MERINGUE, and (iv) spatial distributions representative DE genes. (d, i-iii) All three clusterings also identified a Gfap marked cluster (See Supp. Fig. 17j for corresponding ISH images from the Allen Mouse Brain Atlas, [23, 24]). (iv) Spatial distribution of the Gfap gene.

**Figure 5.**
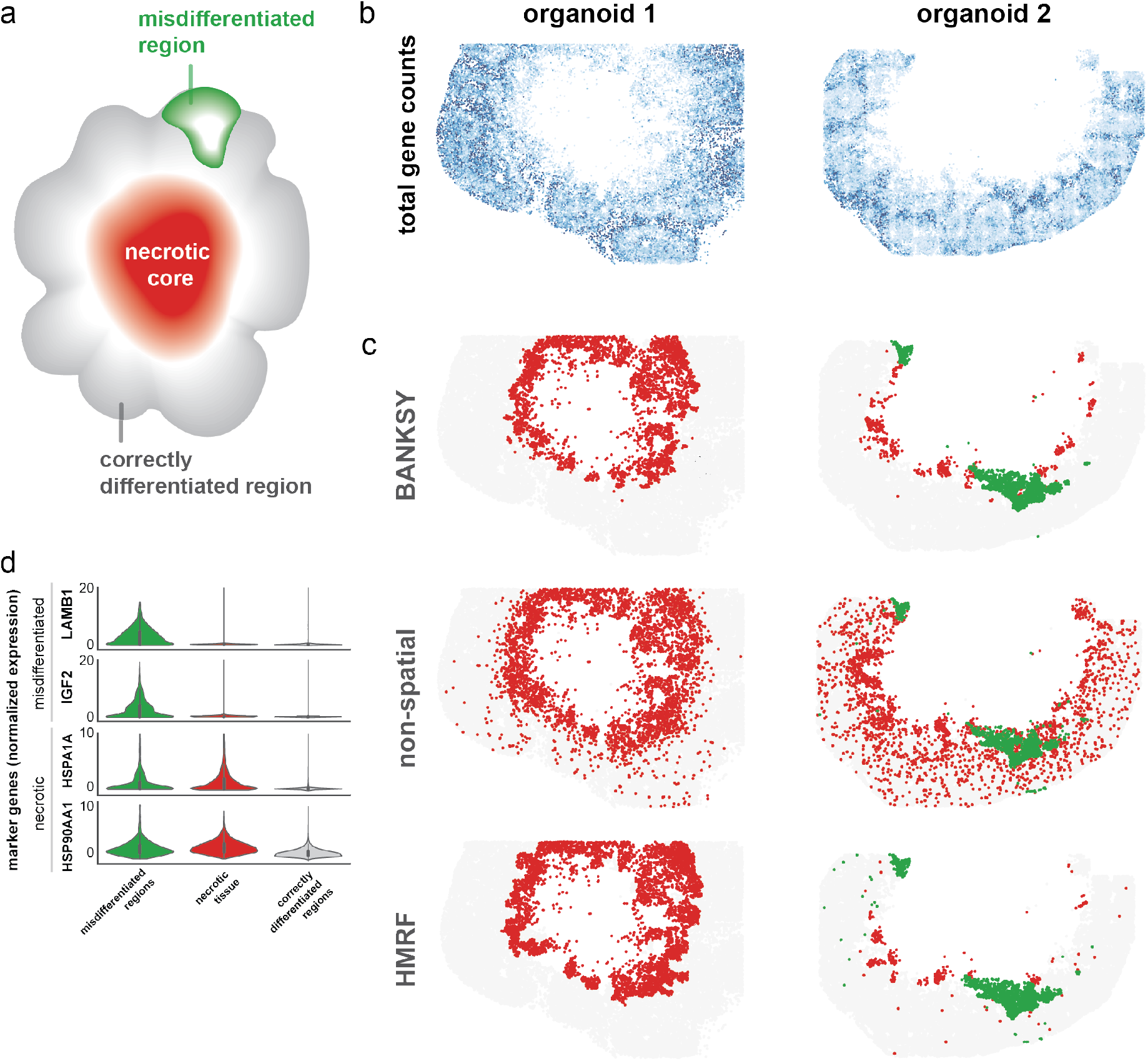
BANKSY’s zone-finding (domain segmentation) mode can be used for performing quality control. (a) Simplified schematic of brain organoid [Lin et al. 2022]. Grey–the correctly differentiated forebrain regions appropriate for the study. Green–a misdifferentiated region. Red–necrotic tissue. (b) Total counts are not necessarily representative of cell type or state, and as such cannot be used to mark out regions of necrotic or misdifferentiated tissue. (c) BANKSY identifies spatially contiguous regions of necrotic and misdifferentiated tissue when applied in zone-finding mode (*λ* = 0.75). Clustering without spatial information and using HMRFs is shown for comparison. We show 2 organoids (out of 3 organoids jointlyprocessed in both BANKSY and non-spatial clustering). HMRF was run on individual organoids; number of clusters set to 10 and *β* set to 0.12 and 0.16 for organoids 1 and 2 respectively. (d) Violin plot of marker genes for the misdifferentiated region and the necrotic core tissue. IGF2 and LAMB1 plots were clipped at a normalised expression level of 20. All normalised expression values were computed using sc-transform.

Instead, we note that one of the principal ways in which a cell’s physical location influences it is through gene expression in its physical microenvironment. Thus, we surmise that the microenvironment signature, and not the raw physical space coordinates or relative locations, are a better medium for encoding spatial information in algorithms. To read out a cell’s microenvironment, BANKSY uses a spatial kernel to compute the weighted average transcriptome of its physical-space neighbours, with cells further away from the index cell having smaller weights (Fig. 1, Methods Section 4.1). We then combine each cell’s microenvironment signature with its transcriptomic features, and present this combined feature set to a graph-based clustering algorithm. To this end, we create a *neighbour-augmented* gene-cell matrix by concatenating ‘self’ and ‘neighbour’ features (Fig. 1a). Assuming *m* measured genes, this concatenation lifts each cell from a *m*-dimensional gene space to a 2*m*-dimensional *neighbour-augmented* gene space, in which cells can be separated by both self expression and microenvironment signature (Fig. 1b, Supp. Fig. 2, Methods Section 4.1). Formally, this space is a Cartesian (or direct) product of two distinct spaces, the first one with coordinates given by the expression of the *m* genes, and the second with coordinates defined by the weighted average expression of these genes in each cell’s physical neighbourhood.

Once the neighbour-augmented matrix has been constructed, it can be fed into any downstream clustering algorithm, in analogy to conventional gene-cell matrices being used with a wide range of algorithms. We have chosen the Leiden community detection algorithm [16] as our default due to its speed and scalability, although we provide integrations with a number of other clustering algorithms (Louvain, mclust, and k-means), in addition to bioinformatics pipelines like Seurat (R, [17]) and Scanpy (Python, [18]).

To control the relative importance of self and neighbour features, we introduce an additional *mixing parameter λ* that adjusts the contribution of neighbourhood features to the total dissimilarity between any pair of cells (Methods, Section 4.1.1, Fig. 1, Sup. Fig. 2). Depending on where *λ* is on the continuum between 0 and 1, BANKSY operates in different qualitative modes (Fig. 2b). When *λ* is small (approximately between 0.1 and 0.3), BANKSY operates in *cell-typing* mode, where the cell’s own transcriptome is dominant relative to its average microenvironment signature. As *λ* increases, the microenvironment exerts a greater influence on the corresponding cluster assignments. In the limit of *λ* approaching 1, the algorithm clusters cells based only on the average microenvironment signature, which leads it to operate in *zone-finding* mode.

To demonstrate that BANKSY is capable of refining cell type assignments and sharpen boundaries between distinct cell layers in data where technical noise may confound cell type assignments, we applied BANKSY in cell-typing mode to the Slide-Seq and Slide-seq V2 data from the mouse cerebellum [3]. Both versions of Slide-Seq offer resolution on the order of a single cell (10 *µ*m), but because the detection beads do not exactly match cell locations, a typical bead contains the transcriptomic signature from one dominant cell with neighbouring cells mixed in. Hence, as noted by the authors of the study, unsupervised clustering performs poorly on such datasets, necessitating deconvolution techniques that rely on scRNA-seq reference data [3, 19]. The reference cell type signatures are used to parse cell type contributions within each bead. We hypothesised that in the absence of reference data, augmenting the partially confounded transcriptomes of individual beads with their microenvironmental signatures would lead to cleaner clustering results. The result of such clustering would be that beads are more accurately assigned to the dominant cell type within the bead.

Applying BANKSY to both the Slide-seq and Slide-seq v2 mouse cerebellum datasets, we found that it more accurately delineated the granular layer, the combined Purkinje neuron and Bergmann glia layer, molecular layer interneurons (MLI) and choroid plexus (Fig. 2a,b, Supp. Figs. 3, 4). In contrast, visual inspection of cell locations from the granular layer cluster showed that unsupervised clustering without spatial information incorrectly classified many more cells outside the layer boundary, consistent with what was reported in [19]. To more quantitatively measure layer cleanness, we computed a normalised connectivity score between each cell type and each other cell type using neighbours defined by the spatial graph connecting each cell to its nearest neighbours (Slide-seq: Fig. 2d, Slide-Seq v2: Supp. Fig. 5). We observed lower cross-connectivity across layers when clustering with BANKSY relative to non-spatial clustering, indicating that the BANKSY clusters were more spatially distinct and less intermingled, confirming our visual assessment.

To assess the accuracy of cell type assignments, we used the reference guided RCTD weights and computed the point-biserial correlation between the major clusters and the RCTD weights for corresponding cell types. We found that in both datasets, the BANKSY clusters had a higher correlation than non-spatial clustering in all clusters except for the oligodendrocyte/polydendrocyte cluster (Supp. Fig. 6). Notably, the astrocyte cluster in Slide-seq v2 showed higher correlation to RCTD despite being interspersed with other cell types.

**Figure 6.**
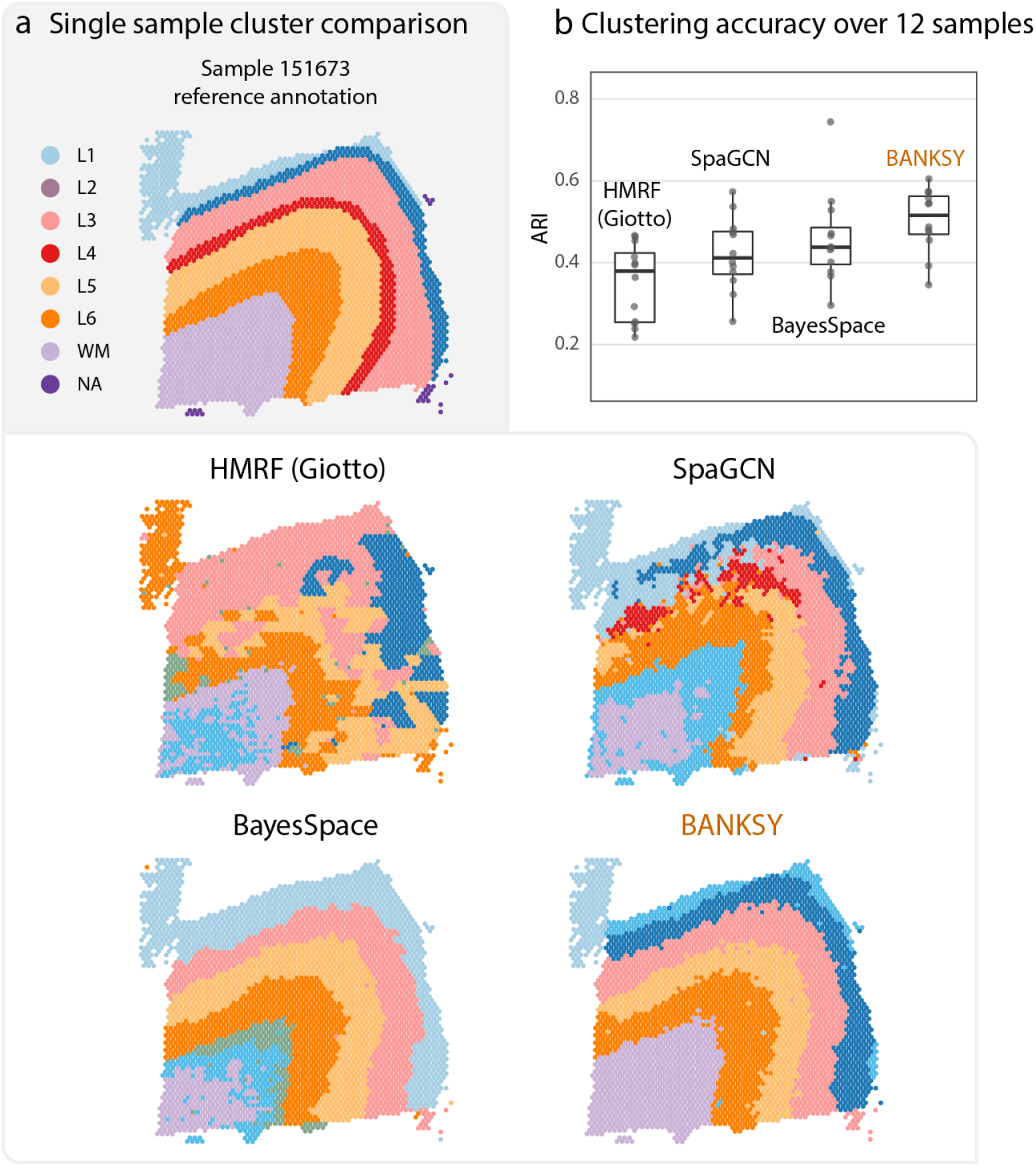
BANKSY’s zone-finding performs better than related methods on the Visium benchmark dataset. (a) Clustering results on sample 151675 of the human DLPFC dataset generated with 10X Visium by BANKSY and selected spatial domain segmentation algorithms. Reference cluster labels from [7] are shown above for comparison. Clusters are coloured by closest match to the manual annotation using the Hungarian algorithm. Clusters not matched to ground-truth labels are arbitrarily coloured. (b) Clustering accuracy comparison for all 12 samples using the Adjusted Rand Index (ARI) metric. Center line, box limits and whiskers represent median, upper and lower quartiles and 1.5*×* inter-quartile range respectively.

To illustrate how BANKSY clustering separates similar transcriptomes using neighbour information, we plotted the averaged expression, or metagene, of the top 20 differentially expressed genes against the equivalent metagene of neighbour expression for two adjacent layers, the granular layer and the Purkinje/Bergmann layer (Fig 2c, Section 3 and Supp. Fig. 8). We observed that while the ‘self’ expression alone had a large overlap between the two layers, cells from each layer were much better separated in the joint self-neighbour (i.e., neighbour-augmented) expression space. A similar effect was observed for other clusters relative to the granular layer (Supp. Figs. 7 and 8).

Next, we hypothesized that in MERFISH datasets, where RNA transcripts are assigned to cells at higher spatial resolution than in Slide-seq, BANKSY may be able to utilize distinct microenvironment signatures to further distinguish different cell subtypes within clusters identified by unsupervised clustering. If present, these cell types would have differing microenvironmental signatures, and perhaps subtly different transcriptomic signatures that are not strong enough to be automatically separated by standard unsupervised clustering. To demonstrate this possibility, we applied BANKSY to a MERFISH mouse hypothalamus dataset spanning a large 3D volume [2]. We first applied non-spatial graph-based clustering to all the cells in the first animal within this dataset, and identified the major clusters found by the authors of that study, including the mature oligodendrocytes and the excitatory neurons (Fig. 3a,b). We found that BANKSY separated the mature oligodendrocytes into two separate clusters that could be matched to a single cluster found in nonspatial unsupervised clustering (Fig. 3c). Mapping these two clusters spatially, we discovered that one cluster corresponded to densely packed mature oligodendrocytes restricted to the anterior commissure of the hypothalamic preoptic region (subcluster 2), and the other to a more diffuse set of mature oligodendrocyte cells spread out throughout the rest of the preoptic region (subcluster 1). Looking at the cells’ own expression signatures within these two subclusters, we found that although the overall gene abundances were almost identical (Supp. Fig. 10i, orange and red clusters), a few genes showed subtle differences in expression between these two subclusters (Supp. Fig. 10ii). Differential expression analysis between these two clusters can be used to identify these genes, and the small differences in their expression between the two subclusters become clearly visible once the expression is z-scaled within only the mature oligodendrocytes subset of cells (Fig. 3d). In particular, the expressions of the Mbp, Lpar1 and Ndrg1 were upregulated in the dense mature oligodendrocyte cluster (subcluster 2) relative to the sparse cluster, and Mlc1, Gad1, Cbln2 and Syt4 were upregulated in the sparser subset of mature oligodendrocytes (subcluster 1). Mbp, Lpar1 and Ndrg1 are involved in the myelination of neurons [20, 21], suggestive of a possible functional difference in myelination by densely packed mature oligodendrocytes in the anterior commissure relative to the sparsely spread out mature oligodendrocytes in the rest of the preoptic area. Plotting the self-neighbourhood metagene signatures (defined in Supp. Section 3), we found that these two oligodendrocyte signatures were easily separable in the neighbour-augmented expression (metagene) space, while showing a large overlap in the self expression space. A similar decomposition into cell-subtypes was also observed for the excitatory neurons (Fig. 3e-h, Supp. Fig. 10iii and 11). This confirmed our hypothesis that BANSKY can distinguish two sub-populations which have very subtly different expression signatures, but otherwise reside in highly distinct microenvironments.

We also tested BANKSY on a mouse hippocampus dataset generated using VeraFISH™, an in-situ barcoded cyclic FISH workflow (Veranome Biosystems, LLC). The results were compared to non-spatial clustering, the spatial clustering module within the MERINGUE pipeline [8] (Fig. 4), and the HMRF module from the Giotto pipeline [22] (Supp. Fig. 12). We found that MERINGUE and non-spatial clustering found very similar results, while BANKSY was able to leverage the distinct neighbourhood signatures to split the clusters into subclusters with subtly distinct transcriptomic profiles (Fig. 4a-c, Supp. Fig. 16a, coloured boxes). For instance, BANKSY separated the cells in the somatosensory cortex from the pyramidal neurons in the CA3 layers (Fig. 4a, i, blue: CA3, light-pink: somatosensory cortex), while non-spatial clustering and MERINGUE did not (Fig. 4a, ii-iii). Examining the spatial distribution of the expression of the top DE genes between these clusters (Supp. Figs. 13, 16) showed that Tbr1, Grm3 and Egr1 were more highly expressed in the somatosensory cortex relative to the CA3 pyramidal neurons (Fig. 4d, i, Supp. Fig. 15) while Cadm3, Parp1 and Clstn2 were more highly expressed in the CA3 neurons. Similarly, BANKSY separated the cells in the fimbria and the thalamic nuclei into two different groups (red and brown clusters, Fig 4b, i) while MERINGUE and non-spatial clustering merged these clusters. These two clusters were distinguished by differential expression of Mobp, Cacna1a, Sgk1, Plp1, and Bcas1 (Fig. 4b, iv, Supp. Figs. 13, 15). BANKSY’s ability to distinguish these clusters stemmed from the fact that the neighbourhood signatures were different between the subclusters in each of these pairs (Supp. Fig. 16a, bottom green and blue boxes in neighbourhood signatures).

The thalamic nuclei region had multiple intermingled clusters of cells (Fig. 4b (brown), c (olive green) and Supp. Fig. 14c, (yellow)). The second of these (olive green) was distinguished from similar cells in the rest of the hippocampus (black) by BANKSY, but not by the other two methods (Fig. 4c, i-iii) and was marked by the expression of Sparcl1 (Supp. Fig. 13iii, 15; general hippocampal area marked by Slc1a3). Taken together, the results for the three pairs of clusters shown in Fig. 4a-c describe an effect similar to what we observed in the mouse hypothalamus dataset in Fig. 3, suggesting that BANKSY can be used to discover subtly different transcriptomic states or cell subtypes on a variety of multiplexed FISH datasets.

Furthermore, for clusters that BANKSY did not split into subclusters, BANKSY mislabelled fewer cells outside the expected cell type regions relative to non-spatial clustering and MERINGUE, similar to what we observed in the Slide-seq cerebellum data. For example, the cluster corresponding to Gfap expression matched the smFISH reference much more closely than equivalent clusters from non-spatial and MERINGUE. (Fig. 4d, i-iv, Gfap marked cluster, and Supp. Fig. 17j showing Allen Mouse Brain Atlas ISH images [23, 24]).

In addition to these three methods, we also tested the performance of the HMRF method from [22], and found that it was not able to distinguish different cell types within single regions, and tended to spatially clump the labels for several clusters (Supp. Figs. 12, 14). Such differences are expected, since this method was designed primarily for zone-finding (domain segmentation), and not cell type identification.

We then applied BANKSY in zone-finding mode for the purpose of spatial domain segmentation. Here, we adjusted lambda such that the average neighbourhood expression signatures were the dominant feature used for clustering cells, to bias the algorithm toward finding zones sharing the same microenvironment. A useful application of this mode is as a selection method for denoting regions of cells to keep or remove in a spatial dataset. We demonstrated BANKSY for this purpose in a companion study [Lin et al. 2022], where initial clustering in zone-finding mode was used to identify regions of misdifferentiated and necrotic tissues (Fig. 5) in slices of brain organoids. Cells in these regions were then removed as part of a preprocessing pipeline for identifying cells following the dorsal forebrain fate. Conversely, non-spatial clustering incorrectly labelled cells within desired regions as necrotic or misdifferentiated. In contrast, BANKSY enabled these regions to be identified and removed cleanly with minimal influence on the regions to be kept (Fig. 5b). We corroborated BANKSY’s zone-finding results using Giotto’s HMRF method [22], which was designed specifically for domain segmentation (Fig. 5c).

Next, we quantitatively tested BANKSY’s spatial domain segmentation capability by applying BANKSY to the LIBD human dorsolateral prefrontal cortex (DLPFC) data generated using the 10x Visium platform [7]. The full dataset consists of 12 slides from 3 subjects, with manual annotations by the authors. These annotations were used as a ground truth comparison to benchmark spatial domain identification in several recent approaches [11, 13, 12]. We benchmarked BANKSY against state-ofthe-art algorithms including BayesSpace [11] and SpaGCN [13] using the adjusted Rand index (ARI) metric proposed by the original authors [7], showing that BANKSY achieved higher median ARI than HMRF (Giotto’s implementation), BayesSpace and SpaGCN, and produced qualitatively good segmentation maps on representative samples 151675 (Fig 6) and 151673 (Supp. Fig. 18). We note that this result was achieved without utilizing histology information for clustering such as in [13]. We also measured clustering accuracy by two other metrics, normalized mutual information distance (NMI) [25] and Matthews correlation coefficient (MCC) [26]. The relative performance of the methods tested was comparable across the three metrics, verifying that BANKSY’s superior performance was not limited to a single metric (Supp. Fig. 19a). In addition, BANKSY is also fast, having shorter run times compared to other methods tested on all 12 datasets, with only [13] having a comparable computational speed (Supp. Fig. 19b).

Finally, we tested how the time required to run BANKSY scales in comparison with other methods as the number of cells in a dataset increases. To this end, we used MERFISH data from Vizgen [27] showing a single coronal slice of the of the mouse brain (Slice 2, Replicate 1, 483 genes, 83463 cells). We cropped the data to five vertical strips (Supp. Fig. 21iii) involving 4k, 8k, 16k, 33k and 83k cells (5 -100% of the data, logarithmically spaced). We then ran conventional (non-spatial) clustering, BANKSY and MERINGUE (all three using Louvain for the clustering step), along with the zone-finding method from Giotto (HMRF, using the expectation-maximisation algorithm) on each of these crops of the data. We found that the total time taken for conventional clustering and BANKSY were similar, HMRFs were between 10-20 times slower, and MERINGUE was between 200-500 times slower (Supp. Fig. 21i, ii). MERINGUE struggled to scale to the full 80k dataset, and had to be terminated once the data size reached about 30 -40k cells due to the algorithm hitting the indexing limits of the R programming language. We note that the BANKSY matrix computation step, which essentially reduces the spatially informed clustering problem to one that can be solved with conventional clustering algorithms (which are used for non-spatial clustering), takes much less time than the clustering step (8.3 seconds vs 94 seconds on the full 83k cells in the dataset; Supp. Fig. 21ii, second table).

## 3 Discussion

In summary, we have developed a principled approach that incorporates information from both a cell’s microenvironment and its own omic profile to generate a combined feature set that is compatible with standard downstream clustering approaches and single-cell analysis pipelines. We have shown that BANKSY is a versatile algorithm that effectively (i) improves cell type assignment for spatially structured cell types, (ii) is capable of distinguishing subtly different cell-subtypes stratified by microenvironment, and (iii) when the microenvironment component within the transcriptomic signature is given sufficient weighting, operates in a *zone-finding* mode, which can be used to identify spatial domains of interest within tissue.

BANKSY introduces a new paradigm for spatial omics analysis that emphasizes microenvironmental features as a salient feature for cell type prediction. Rather than relying on the common assumption that a cell is likely to be of the same type as its neighbours [8, 11, 10], BANKSY retains the cells’ unique omic signature while using its microenvironment to augment predictions of cell type.

The key feature of BANKSY that enables cell-typing is the explicit separation of self and neighbour transcriptomes. Summing this microenvironment signature with the cell’s own omic profile in highly heterogeneous tissues is problematic from a cell-typing perspective. This is because it averages out the profiles of unique cells embedded in a uniform environment of another cell type, an extreme example being infiltrating immune cells in a tumour. In practice, we found that this merely smoothed out cluster assignments of cells, leading to spatially homogeneous clusters that hide finer biological structures.

Such an approach is ideal for discovery of spatial domains in tissue, and has been used effectively in a variety of recent approaches [12, 13]. However, when cell-typing is the goal, this may lead to poorer resolution and accuracy, particularly in heterogeneous regions of tissue with mixtures of cell types.

Recent deep learning approaches [13] based on graph convolutional networks also implicitly utilize microenvironment information by aggregating neighbour features. However, this architecture *sums* self and neighbour features, merging them into a single feature-set, making this approach more suited for spatial domain segmentation. In addition, feature transformations may reduce the interpretability of the data, and require additional training and parameter optimization steps.

With a slight adjustment of parameters, BANKSY is also capable of detecting spatial domains, as shown on the brain organoid and DLPFC Visium data. For data types where each location represents a single cell, such as the brain organoid multiplexed FISH data, raising lambda weights the local microenvironment over cell state allowing for detection of spatial zones. For data types where each spot covers multiple cells and hence already represents an averaged transcriptomic signature, BANKSY will find spatial domains at lower lambda values. We showed this on the DLPFC Visium dataset, where BANKSY’s performance exceeded that of recently published spatial domain identification algorithms [11, 13], showing the versatility and applicability of the BANKSY framework to the important goal of spatial domain segmentation. Hence, BANKSY offers a unified and computationally tractable approach for both cell type identification and domain segmentation.

BANKSY is broadly applicable to a wide variety of spatial omic data types from a diverse set of technologies. We have demonstrated that BANKSY greatly improves analysis of spatial transcriptomic data both in sequencing based technologies and in multiplexed FISH based approaches. In principle, BANKSY is applicable to any spatial omics technology, including proteomics methods that allow measurement of omics profiles at the single cell resolution, and future work will include applying BANKSY to new technologies that measure other aspects of cell state. In addition, BANKSY may be easily extended to use combinations of feature types in multiomic datatypes. For example, most Visium datasets include accompanying histology images which some existing methods utilize to augment transcriptomic profiles [13, 12]. Including appropriate image features (represented by colour channel data [13] or features extracted using pre-trained neural networks [12]) in combination with gene expression may further improve accuracy of cell-typing or spatial-domain identification.

We showed that BANKSY has practical implications for the analysis of emerging spatial omics technologies. It enabled unsupervised clustering to be performed on Slide-Seq data, which was previously infeasible, allowing users to perform analysis on such data where corresponding reference data are not available. Compared to the reference-based approach, BANKSY has the advantage of being unbiased, avoiding platform effects inherent to cell type imputation, and may allow for detection of cell types absent from reference data. However, BANKSY is not a replacement for reference-based methods as, like any unsupervised clustering approach, it cannot detect multiple cell types in a single bead, and does not improve on non-spatial clustering when separating cell types sharing the same microenvironment, such as with intermingled Purkinje neurons and Bergmann glia in the cerebellum. Hence, BANKSY provides a complementary alternative to reference-based supervised approaches, and will broaden the range of insights that can be obtained from sequencing-based spatial omics methods. With the brain organoid MERFISH dataset, we showed that BANKSY’s zone-finding mode is useful for quality control, distinguishing broad regions of tissue with desired or undesired characteristics, and hence is a useful addition to the toolbox for analysing multiplexed FISH data.

Given any traditional single-cell analysis algorithm that uses a gene-cell matrix, one can posit that replacing its gene-cell matrix with BANKSY’s neighbour-agumented version transforms it from a non-spatial algorithm to a spatially-informed one. This position leverages the underlying insight that the microenvironment expression is precisely what mediates any influences on a cell that may exist due to differing spatial locations, and is therefore the correct medium to encode spatial information in algorithms. Indeed, throughout this study, we have demonstrated that in the context of clustering, there is significant value-add to this approach for both cell-typing and zone-finding. Another context where this could be tested is for identifying spatially variable genes. For instance, clustering could be performed in zone-finding mode, and then the DE-genes between these spatially contiguous zones would be the spatially variable genes, an approach that was used effectively in [13]. This could be used as an unbiased alternative to model-based methods like SpatialDE and SPARK [28, 29], which assume, for instance, that the gene expression is correlated in physical space via various kinds of kernels.

BANKSY is computationally efficient and scales well to large datasets. The computational complexity of the neighbour matrix computation increases linearly with the number of cells, given a fixed number of genes and cell neighbours, making it a feasible approach even on much larger datasets. In fact, we found that the clustering step (Supp. Fig. 21ii, second table) dominated the computation time in most datasets we tested, hence total computation times for BANKSY were comparable to non-spatial clustering on all dataset sizes tested (4k cells to 80k cells). In practice, we found that BANKSY and conventional clustering were tens to hundreds of times faster than other methods on typically sized datasets (Supp. Fig. 21).

In addition, BANKSY easily integrates into existing single-cell analysis pipelines, allowing for the application of feature selection tools before, and visualisation tools after its use. We provide an implementation of the algorithm as a standalone R package, a plug-in to the Seurat pipeline [17] and to the SinceCellExperiment framework [30], as well as a stand-alone Python version that is compatible with the ScanPy package [18].

All in all, BANKSY’s biologically inspired approach offers an accurate, sensitive, and scalable spatial clustering tool that unifies cell type identification and domain segmentation into a single framework.

## 4 Methods

### 4.1 The BANKSY algorithm

In this section we describe the BANSKY algorithm in the context of clustering a set of cells, based on the expressions of a set of genes and the locations of the cells or capture-spots (used in sequencing-based technologies) in physical space. Briefly, we construct a neighbourhood graph between the cells in physical space, and use this to compute the (weighted) average neighbourhood gene expression vector for each cell. This vector is concatenated with the original expression vector associated to this cell to give the coordinates of the cell in a joint own expression-neighbourhood expression (product) space, which we refer to as the neighbour-augmented space. In what follows, we describe this procedure formally.

Let a set of cells arranged in physical space be indexed by the set ℐ = {1, 2, …, *N*}, and have a set of spatial coordinates 𝒳= *x*_*i*_ ℝ^*d*^ | *i*∈ ℐ, where *d* is typically 2 or 3. For each cell, assume that the expression of the same set of *m ∈ℤ* _*≥* 1_ genes has been measured, so that the expression information can be expressed as a gene-cell expression matrix *C* = [*c*_1_ *c*_2_ … *c*_*N*_] ℝ ^*m×N*^, where *c*_*i*_ ∈ ℝ ^*m*^ is the expression of the *m* genes in cell *i*.

Next, assign a set of neighbours to each cell on the basis of their relative locations in physical space. That is, associate with the *i*-th cell a neighbourhood,

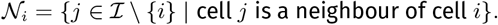

Example of schemes to choose neighbours can be Delaunay triangulation, K-nearest neighbours, or all neighbours encompassed by a disc of fixed radius. Once chosen, the set of neighbourhood relationships can be encoded in a directed graph in physical space, where each cell is a vertex, and there is an edge from cell *j* to cell *i* if *j∈ 𝒩* _*i*_. We use k-nearest neighbours as our default to define the neighbourhood, with *k* = 10.

We also find it useful to associate weights with each edge, which can be used to allow nearby cells to contribute more to a cell’s neighbourhood signature relative to cells further away. The weighting can be done using some monotonically non-increasing function of the distance between the cells connected by each edge. Choices of such kernel functions include 1*/r* or the Gaussian kernel, 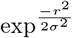, where *r* is the (Euclidean) distance between the pair of cells under consideration, and *σ* is a parameter that controls how large the effective size of the neighbourhood is. Other kernels can be a simple uniform kernel (no weighting) or what we call the rank-Gaussian kernel, with a weight of 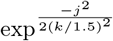 for the *j*-th neighbour (in physical space), up to a fixed maximum number of *k* neighbours, and *k/*1.5 controlling how quickly the drop-off in influence occurs with the rank of the neighbour. While *k* is fixed when the neighbourhoods are chosen using K-nearest neighbours, using a Delaunay triangulation or considering all cells within a disk as neighbours leads to a different number of neighbours per cell, and for the *i*-th cell, one may replace the constant *k* with an *i*-dependent *k*_*i*_ =| 𝒩 _*i*_ | in the above expression for the rank-Gaussian kernel (| · | denoting the cardinality of the set of neighbours).

As a default, we use a 1*/r* kernel, although we find that in practice the results are insensitive to variations in these choices (Supp. Fig. 20iii). The 1*/r* kernel was chosen to down-weight cells by distance, especially in cases where a subset of the *k* = 10 cells are much further that the remaining cells. In addition, the sum of cell-neighbour weights are normalised to 1, which leads to similar weightings in regions of varying density. Let *w*_*ij*_ *∈* ℝ denote the weight associated with the edge from cell *j* to index cell *i*, and *r*_*ij*_ *∈* ℝ be the physical distance between them (in any units, since *w*_*ij*_ below is dimensionless). Then,

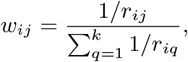

where the denominator ensures that the weights associated with a cell’s neighbourhood sum to one. The normalisation with other weighting kernels is done analogously. With this set up, we can compute the average neighbourhood signature associated with the *i*-th cell as

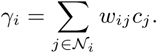

We concatenate these neighbourhood expression vectors into a neighbour expression gene-cell matrix,

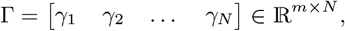

and concatenate *C* and Γ into the *neighbour-augmented matrix* or (*BANKSY matrix*),

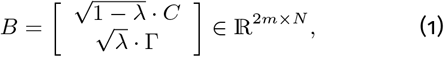

where *λ* ϵ [0, 1] weights the relative contributions of the two component matrices.

This matrix can now be used as a feature-cell matrix for any clustering algorithm, such as hierarchical clustering or graph-based clustering. Due to the large number of cells involved in most genomics studies, graph-based clustering methods such as the Louvain or Leiden algorithms that have gained popularity in recent years, and work well with feature-object matrices of the type in equation (1). In this study, we use the Leiden clustering algorithm, although Louvain clustering or even k-means clustering work well too, and can have even faster run times in practice.

#### 4.1.1 Convex Combinations of Distance Matrices

In this section, we show that the weighted concatenation of feature-object matrices that BANKSY takes is equivalent to the overall cell-cell distance matrix being a convex combination of the distance matrices corresponding the the component feature object matrices,

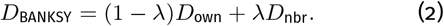

This illuminates an alternate point of view, whereby we see that taking convex combinations of dissimilarity matrices is a general framework for combining different sources of dissimilarity between objects, or of adding ‘soft constraints’ [31] to clustering problems. In the BANKSY framework, we are effectively adding a neighbourhood dissimilarity term to the traditional transcriptomic dissimilarity used in conventional non-spatial clustering. In principle, other sources of dissimilarity may be included as well.

Let *C*, Γ and *B* be as in Section 4.1, and recall that the *i*^th^ column of the BANKSY matrix *B* is

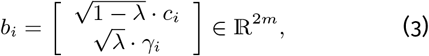

where the dot (·) simply denotes multiplication of a scalar and a vector, and *m* is the number of genes.

Define the Euclidean distance matrix corresponding to *C*,

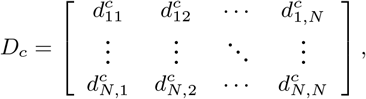

with 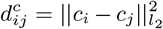 defining the (squares of) the *l*_2_ distances between cells. The neighbourhood and BANKSY distance matrices, *D*_*γ*_ and *D*_*b*_, are analogously defined. With these definitions, it follows immediately (via the Pythagoras’ theorem on Equation (3)) that *D*_*b*_ = (1 − *λ*)*D*_*c*_ + *λD*_*γ*_, which is just Equation (2).

### 4.2 Software and Data Availability

The R package can be obtained from https://github.com/prabhakarlab/Banksy, while the python version is available from https://github.com/prabhakarlab/Banksy_py.

For the SlideSeq mouse cerebellum dataset, the data were obtained from https://github.com/RubD/spatial-datasets/blob/master/data/2019_slideseq_cerebellum/raw_data/slideseq_cerebellum_urls.txt. We used the BeadLocationsForR and MappedDGEForR

.csv files, filtering for total gene count and minimum genes detected. For Slide-seq V2, we obtained the data from the Broad Institute Single Cell Portal at https://singlecell.broadinstitute.org/single_cell/study/SCP948.

The MERFISH mouse hypothalamus data were obtained from https://doi.org/10.5061/dryad.8t8s248,

The mouse hippocampus data were collected using the VeraFISH assay (Veranome Biosystems, LLC, Mountain View, CA, USA) as described in Supp. Section 2.2. The data are available via the R GitHub page.

The human brain organoid data were collected inhouse and obtained as specified in Lin et al. (2022).

The 10x Visium data of the dorsolateral prefrontal cortex (DLPFC) was obtained from the spatialLIBD project (http://spatial.libd.org/spatialLIBD).

## 5 Author Contributions

BANKSY was conceived by VS and SP, with help from HKL and KHC, and with major conceptual contributions in its further development and demonstrations by NC and JLee. JLee and VS designed and wrote the R package, and NC built the Python implementation. JLiu contributed to the design of the application to brain organoid data. The organoid and VeraFISH data were provided by JLiu, WKC, LL, YCC and ET. VS, NC, JLee, SP and KHC wrote the manuscript, and all authors edited it.

## Acknowledgements

We are grateful to Anandh Swaminathan, Bobby Ranjan, Grace Yeo, Xinrui Zhou and Reema Baskar for useful discussions, and to Jen Yi Wong for help with Linux administration and scripting.

## Notes

https://github.com/prabhakarlab/Banksy

### Competing Interest Statement

ET and YCC are employed by Veranome Biosystems.

## References

[1] Kok Hao Chen et al. “Spatially resolved, highly multiplexed RNA profiling in single cells”. In: Science 348.6233 (2015). issn: 0036-8075. doi: 10.1126/science.aaa6090.

[2] Jeffrey R. Moffitt et al. “Molecular, spatial, and functional single-cell profiling of the hypothalamic preoptic region”. In: Science 362.6416 (Nov. 2018). issn: 0036-8075, 1095-9203. doi: 10.1126/science.aau5324. (Visited on 03/07/2021).

[3] Samuel G. Rodriques et al. “Slide-seq: A scalable technology for measuring genome-wide expression at high spatial resolution”. In: Science 363.6434 (2019), pp. 1463–1467. issn: 0036-8075. doi: 10.1126/science.aaw1219.

[4] Tongtong Zhao et al. “Spatial genomics enables multi-modal study of clonal heterogeneity in tissues”. In: Nature 601.7891 (Jan. 2022), pp. 85–91. issn: 1476-4687. doi: 10.1038/s41586-021-04217-4.

[5] Felix J. Hartmann et al. “Single-cell metabolic profiling of human cytotoxic T cells”. In: Nature Biotechnology 39.2 (Feb. 2021), pp. 186– 197. issn: 1546-1696. doi: 10.1038/s41587-020-0651-8.

[6] Chee-Huat Linus Eng et al. “Transcriptome-scale super-resolved imaging in tissues by RNA seqFISH+”. en. In: Nature 568.7751 (Apr. 2019), pp. 235–239. issn: 1476-4687. doi: 10.1038/s41586-019-1049-y.

[7] Kristen R. Maynard et al. “Transcriptome-scale spatial gene expression in the human dorsolateral prefrontal cortex”. In: Nature Neuroscience 24.3 (Mar. 2021), pp. 425–436.

[8] Brendan F Miller et al. “Characterizing spatial gene expression heterogeneity in spatially resolved single-cell transcriptomics data with nonuniform cellular densities”. In: Genome Research (2021). doi: 10.1101/gr.271288.120.

[9] Haotian Teng, Ye Yuan, and Ziv Bar-Joseph. “Clustering spatial transcriptomics data”. In: Bioinformatics 38.4 (Feb. 2022), pp. 997– 1004. issn: 1367-4803. doi: 10.1093/bioinformatics/btab704.

[10] Qian Zhu et al. “Identification of spatially associated subpopulations by combining scRNAseq and sequential fluorescence in situ hybridization data”. In: Nature biotechnology (Oct. 2018), 10.1038/nbt.4260. issn: 1546-1696. doi: 10.1038/nbt.4260.

[11] Edward Zhao et al. “Spatial transcriptomics at subspot resolution with BayesSpace”. In: Nature Biotechnology 39.11 (Nov. 2021), pp. 1375– 1384. issn: 1546-1696. doi: 10.1038/s41587-021-00935-2.

[12] Duy Pham et al. “stLearn: integrating spatial location, tissue morphology and gene expression to find cell types, cell-cell interactions and spatial trajectories within undissociated tissues”. In: bioRxiv (2020). doi: 10.1101/2020.05.31.125658.

[13] Jian Hu et al. “SpaGCN: Integrating gene expression, spatial location and histology to identify spatial domains and spatially variable genes by graph convolutional network”. In: Nature Methods 18.11 (Nov. 2021), pp. 1342– 1351. issn: 1548-7105. doi: 10.1038/s41592-021-01255-8.

[14] Vincent D. Blondel, Jean-Loup Guillaume, Renaud Lambiotte, and Etienne Lefebvre. “Fast unfolding of communities in large networks”. en. In: Journal of Statistical Mechanics: Theory and Experiment 2008.10 (Oct. 2008), P10008. issn: 1742-5468. doi: 10.1088/1742-5468/2008/10/P10008.

[15] V. A. Traag, L. Waltman, and N. J. van Eck. “From Louvain to Leiden: guaranteeing well-connected communities”. In: Scientific Reports 9.1 (Mar. 2019), p. 5233. issn: 2045-2322. doi: 10.1038/s41598-019-41695-z.

[16] V. A. Traag, L. Waltman, and N. J. van Eck. “From Louvain to Leiden: guaranteeing well-connected communities”. In: Scientific Reports 9.1 (Mar. 2019), p. 5233. issn: 2045-2322. doi: 10.1038/s41598-019-41695-z.

[17] Tim Stuart et al. “Comprehensive Integration of Single-Cell Data”. In: Cell 177.7 (June 2019), 1888–1902.e21. issn: 0092-8674. doi: 10.1016/j.cell.2019.05.031.

[18] F. Alexander Wolf, Philipp Angerer, and Fabian J. Theis. “SCANPY: large-scale single-cell gene expression data analysis”. In: Genome Biology 19.1 (Feb. 2018), p. 15. issn: 1474-760X. doi: 10.1186/s13059-017-1382-0.

[19] Dylan M. Cable et al. “Robust decomposition of cell type mixtures in spatial transcriptomics”. In: Nature Biotechnology (Feb. 2021). issn: 1546-1696. doi: 10.1038/s41587-021-00830-w.

[20] Dongqiong Xiao et al. “The Roles of Lpar1 in Central Nervous System Disorders and Diseases”. In: Frontiers in Neuroscience 15 (2021). issn: 1662-453X. doi: 10.3389/fnins.2021.710473. (Visited on 04/06/2022).

[21] Damien Marechal et al. “N-myc downstream regulated family member 1 (NDRG1) is enriched in myelinating oligodendrocytes and impacts myelin degradation in response to demyelination”. In: Glia 70.2 (2022), pp. 321– 336. issn: 1098-1136. doi: 10.1002/glia.24108.

[22] Ruben Dries et al. “Giotto: a toolbox for integrative analysis and visualization of spatial expression data”. In: Genome Biology 22.1 (Mar. 2021), p. 78. issn: 1474-760X. doi:10.1186/s13059-021-02286-2.

[23] Ed S. Lein et al. “Genome-wide atlas of gene expression in the adult mouse brain”. In: Nature 445.7124 (Jan. 2007), pp. 168–176. issn: 1476-4687. doi: 10.1038/nature05453. (Visited on 04/11/2022).

[24] Allen Mouse Brain Atlas [dataset]. 2011. url: http://mouse.brain-map.org.

[25] Nguyen Xuan Vinh, Julien Epps, and James Bailey. “Information Theoretic Measures for Clusterings Comparison: Variants, Properties, Normalization and Correction for Chance”. In: J. Mach. Learn. Res. 11 (Dec. 2010), pp. 2837– 2854. issn: 1532-4435.

[26] Davide Chicco, Niklas Tötsch, and Giuseppe Jurman. “The Matthews correlation coefficient (MCC) is more reliable than balanced accuracy, bookmaker informedness, and markedness in two-class confusion matrix evaluation”. In: BioData Mining 14.1 (Feb. 2021), p. 13. issn: 1756-0381. doi: 10.1186/s13040-021-00244-z.

[27] Vizgen Data Release Program. May 2021. url: https://vizgen.com/support/data-release-program.

[28] Shiquan Sun, Jiaqiang Zhu, and Xiang Zhou. “Statistical analysis of spatial expression patterns for spatially resolved transcriptomic studies”. In: Nature Methods 17.2 (Feb. 2020), pp. 193–200.

[29] Valentine Svensson, Sarah A. Teichmann, and Oliver Stegle. “SpatialDE: identification of spatially variable genes”. In: Nature Methods 15.5 (May 2018), pp. 343–346.

[30] Robert A. Amezquita et al. “Orchestrating single-cell analysis with Bioconductor”. en. In: Nature Methods 17.2 (Feb. 2020), pp. 137– 145. issn: 1548-7105. doi: 10.1038/s41592-019-0654-x.

[31] Marie Chavent, Vanessa Kuentz-Simonet, Amaury Labenne, and Jérôme Saracco. “ClustGeo: an R package for hierarchical clustering with spatial constraints”. In: Computational Statistics 33.4 (Dec. 2018). 1707.03897, pp. 1799–1822. issn: 0943-4062, 1613-9658. doi: 10.1007/s00180-018-0791-1.

