## supplemental text and figures for "BANKSY: A Spatial Omics Algorithm that Unifies Cell Type Clustering and Tissue Domain Segmentation"

April 11, 2022

### 1 Directly Appending Spatial Coordinates

As mentioned in the Results section, the simple approach of incorporating spatial information into clustering methods is to append the spatial coordinates of cells to their gene expression vectors.

As in Section 4.1, let the cells be indexed by the set $\mathcal{I} = \{1, 2, \ldots, N\}$, with spatial coordinates $\mathcal{X} = \{x_i \in \mathbb{R}^d \mid i \in \mathcal{I}\}$, where $d$ is typically 2 or 3. Let us collect the coordinates of the centroids of cells into a matrix,

$$X = \left[ \begin{array}{cccc} x_1 & x_2 & \ldots & x_N \end{array} \right] \in \mathbb{R}^{d \times N}, \tag{1}$$

and as before, we define the gene-cell expression matrix by $C = \left[ \begin{smallmatrix} c_1 & c_2 & \cdots & c_N \end{smallmatrix} \right] \in \mathbb{R}^{m \times N}$, where $c_i \in \mathbb{R}^m$ is the expression of the $m$ genes in cell $i$.

In this approach, we create an augmented feature-cell matrix,

$$A = \left[ \begin{array}{c} \sqrt{1-\lambda} \cdot C \\ \sqrt{\lambda} \cdot X \end{array} \right] \in \mathbb{R}^{(m+2) \times N}, \tag{2}$$

which, in terms of distance matrices, is just $D_{\text{direct}} = \lambda D_{\text{expression}} + (1-\lambda) D_{\text{physical}}$, where $D_{\text{expression}}$ is the matrix of cell-cell distances in expression space, and $D_{\text{physical}}$ is the matrix of pairwise distances in physical space, $\lambda$ is a mixing parameter that controls the relative weighting of these two types of distances, and $D_{\text{direct}}$ is the resulting 'spatially informed' distance matrix in this approach (corresponding to feature-cell matrix A in Equation (2)).

This is similar to the approach taken in the geo-spatial literature [3], where adjacent municipalities in southern France had to be grouped together based on their socio-economic features (analogous to genes in our data), subject to a soft constraint that adjacent municipalities be labelled with the same label.

For spatial omics, this approach does not work because it tends to group cells that are physically near each other into the same cluster, and label physically distant cells differently (Supp. Fig. 1). This is antithetical both to cells of multiple different cell types being interspersed within a single region and to a single cell type being located in different regions of tissue (an example of the latter being, for instance, the two hemispheres of the brain, Supp. Fig. 1b). For large regions, it can even break them up into smaller 'patches', as seen for the cortical layers in Supp. Fig. 1c, d.

### 2 Data Analysis Pipelines

#### 2.1 Mouse Cerebellum Slide-Seq data

For the Slide-seq cerebellum dataset, we filtered out cells with less than 20 and more than 1000 gene counts. From Slide-seq v2, we filtered out cells with less than 50 and more than 2500 gene counts. For both datasets, we removed genes present in less than 10 cells, and used Scanpy's implementation of Seurat's highly-variable gene (HVG) function on log-transformed normalised counts to select the top 2000 HVGs. Log-transformation was only used for HVG computation and was not used for the subsequent steps of the BANKSY algorithm. For Slide-seq v2, we also removed all bead locations beyond a fixed radius of 2550 $\mu m$ around an approximate center point of the puck.

We then normalised total gene counts per cell to the median gene count. After performing the neighbour matrix computation, we z-scaled each row of the self and neighbour matrices (i.e., centered the means, and scaled the standard deviations of each feature to 1, across all cells). The resulting scaled matrices were combined into the neighbour-augmented (BANKSY) matrix with $\lambda = 0.2$.

The physical-space neighbourhood graph was built using the bead locations, with a k-nearest neighbours approach ($k_{\text{geom}} = 10$) and a $1/r$ kernel to weight the edges of the graph, as described in Section 4.1. The sum of weights across a cell's neighbours was normalised to 1.

Next, we used principal component analysis (PCA) to reduce the dimensionality of the neighbour-augmented matrix to 16 principal components, and used the Leiden clustering [12] algorithm with resolution set to $r = 0.6$, and the number of expression space neighbours set to $k_{\text{expr}} = 30$. For 'non-spatial clustering', we performed the same procedure on the original normalised and z-scored gene-cell matrix, with the same number of PCs, the same clustering algorithm, and the same resolution parameters. Further optimisation of parameters (resolution and number of PCs) for non-spatial clustering did not noticeably improve clustering results relative to BANKSY clustering. The cluster labels from non-spatial analysis were matched to corresponding clusters from BANKSY using the Hungarian algorithm, as described in Section 4, with one exception: the non-spatial cluster for molecular layer interneurons (MLI) was merged with a group of low quality spots at the edge of the puck, hence incorrectly matched to a BANKSY cluster containing only these low-quality spots. The cluster that was originally matched to BANKSY's MLI cluster had very low correspondence (r of 0.01) to the reference cell-type signature as estimated by RCTD. Hence we manually reassigned the MLI cluster to the more appropriate cluster for non-spatial clustering. From the dimension reduction step onward, the procedure and parameters used were identical for both Slide-seq and Slide-seq v2.

We then manually matched clusters to the closest reference cluster visualised using RCTD weights and the metagene (defined in Supp. Section 3) was computed from top DE genes in the reference scRNA seq dataset (dropviz.org). Subsequent correlation analysis (Sup. Fig. 6) confirmed that the clusters were correctly matched. Where appropriate, clusters were merged, as was done in [2] for the granular layer. In Slide-seq, the granular layer, MLI, and oligodendrocytes/polydendrocytes each consisted of 2 clusters which were merged into a single cluster. In Slide-seq v2, BANKSY clustering yielded 2 clusters for the granular layer which were merged, while non-spatial clustering yielded a single cluster that was matched to one of the BANKSY clusters. The separate clusters are shown in Sup. Figs. 3 and 4.

#### 2.2 Mouse Hippocampus VeraFISH Data

Six weeks old, female, C57BL/6NTac mice were purchased from InVivos, Singapore. They were euthanised and dissected for removal of the brain from the animals. All animal procedures were done in accordance with the approved Institutional Animal Care and Use Committee (IACUC) protocol (Protocol #211580) obtained from the IACUC of the biomedical resource center. Mouse brains were embedded in optimal cutting temperature compound (Sakura, USA), frozen and stored at -80 °C.

Veranome Biosystems' VeraFISH assay was used for data collection. Tissue samples were permeabilised using 70% ethanol overnight and incubated with VeraFISH sample prep wash buffer for 2 h at 47 °C. After the overnight incubation in VeraFISH Target Probe and Hyb Buffer, samples were rinsed with VeraFISH sample prep wash buffer for 1 h at 47 °C and stored with the VeraFISH Cycling Buffer at 4 °C till imaging. Tissue samples were imaged in fully automatic VSA-1 Imager from Veranome Biosystems that includes incubation with VeraFISH Barcode Probes and imaging through multiple hybridisation processes. The multiplexed images were then processed through VeraWorks© software to reconstruct the spatial coordinates of VeraFISH Barcode Probes. A Mask-RCNN based deep learning pipeline was used to segment DAPI stained nuclei and generate single cell expression matrix in .csv files (`cellmatrix.csv`).

Once we had the raw gene counts per cell, we removed cells with total counts less than the 5-th percentile, and greater than the 98-th percentile. We also removed any genes with expression (at least one count) in less than 1% of cells. The total counts in each cell in the resulting gene-cell matrix were normalised to a value of 100, and the neighbour expression matrix was computed with $k_{\text{geom}} = 10$ and the default $1/r$ weighting kernel. We then z-scaled the rows of both the own expression and neighbour expression matrices, and combined them into the full neighbour-augmented (BANKSY) matrix at two values of lambda (0 and 0.3). Finally, we performed PCA, keeping the top 20 PCs, and clustered using Leiden clustering with a resolution of 1.2 and $k_{\text{expr}} = 50$.

For the MERINGUE runs on this dataset, we used the standard pipeline from [9]. Briefly, MERINGUE computes a Delaunay triangulation graph between the physical locations of cells, with a maximum threshold distance within which cells were considered neighbours (given by the `filterDists` argument in the `getSpatialNeighbors` function; we chose a conservative threshold of 750 pixels, such that most adjacent cells were defined to be neighbours). It then computes the shortest path-length (geodesic) distance between each pair of cells that are neighbours in the transcriptome space graph. These distances are used to weight the edges of the transcriptome space graph during graph-based clustering. We used 20 PCs and $k_{\text{expr}} = 15$ (MERINGUE) and $50$ (MERINGUE low resolution, see next paragraph) in MERINGUE's spatial clustering function, `getSpatiallyInformedClusters`.

In Fig. 4, non-spatial clustering and BANKSY had two parameters that controlled the effective number of clusters (or the effective clustering 'resolution'): the $k_{\text{expr}}$ and `resolution` parameters. The values used for these parameters were $k_{\text{expr}} = 50$ and `resolution` = 1.2. The MERINGUE pipeline, on the other hand, only had a $k_{\text{expr}}$ parameter, which presented us with the choice between picking the same parameter value ($k_{\text{expr}} = 50$) or tuning its value until the *results* were similar to those from non-spatial clustering and BANKSY (so that the 'effective' resolution would be that of the non-spatial clustering and BANKSY). For completeness, we have shown the results for both cases. We believe that because the parameters between MERINGUE and non-spatial clustering do not have a one-to-one correspondence, the 'effective' resolution method of comparing the runs is better grounded, and therefore have included this as the primary comparison in Fig. 4, while the run with $k_{\text{expr}} = 50$ is shown in the supplementary figures as 'MERINGUE (low resolution)'. The value of $k_{\text{expr}}$ for the MERINGUE run shown in the main figure is 15.

#### 2.3 Mouse Hypothalamus MERFISH Data

The data for the mouse hypothalamus atlas study [10] were downloaded from the link provided in Data Sources (Section 4.2). We used the data for Animal 1, which comprised 12 z-slices, 50 $\mu$m apart, labelled Bregmas -0.29 mm to +0.26 mm in increments of 0.05 mm by the authors of that study (and interchangeably labelled -290 $\mu$m, -240 $\mu$m, $\ldots$ +260 $\mu$m elsewhere in their data, and in Supp. Fig. 9). The gene Fos was removed from the dataset, because it contained 'NaN' (Not a Number) entries. This dataset had the gene counts normalised by the imaged volume of each cell, and we normalised each column's total expression to 100 before using the resulting gene-cell matrix to compute the BANKSY matrix at $\lambda \in \{0, 0.2\}$. The physical-space neighbourhood graph was built using the cell locations, with a k-nearest neighbours approach ($k_{\text{geom}} = 15$) and a $1/r$ kernel to weight the edges of the graph, as described in Section 4.1. We then z-scaled the means and standard deviations of each row (feature) of the BANKSY matrix to 0 and 1 respectively. Conceptually, it seems preferable to perform the z-scaling step after the average neighbourhood expression computation, as this ensures that the variance of the own and neighbour expression features in the neighbour-augmented matrix are equal. In practice, however, we find that z-scaling before average neighbourhood expression computation produces similar clustering results.

Next, we used principal component analysis to reduce the dimensionality of the BANKSY matrix to 20 principal components, and clustered the cells using the Leiden algorithm [12] with resolution $r = 0.5$, and number of transcriptome space neighbours $k_{\text{expr}} = 50$. The resulting cluster labels were harmonised by solving the linear sum assignment problem using the Hungarian algorithm, as described in Supp. Section 4.

#### 2.4 Brain Organoid MERFISH Data

The data were obtained as specified in [Lin et al. 2022]. For this analysis, we used three control datasets. Normalised counts were obtained using sc-transform [11], and genes were filtered as described in [Lin et.

al 2022]. After computing the neighbour-augmented matrices for each dataset individually we z-scaled all rows (features) of each neighbour augmented matrix individually, and then concatenated the matrices along the columns (cells) to enable use of all available cells for clustering. We then performed dimension reduction using PCA, keeping the first 15 PCs for clustering analysis. The parameters used were $k_{\text{geom}} = 10$ and $\lambda = 0.75$, with spatial neighbourhoods defined using k nearest neighbours. Finally, we performed Leiden clustering with resolution = 1.0 and $k_{\text{expr}} = 150$. For HMRF comparison, we used the implementation in Giotto 1.1.0, and performed clustering separately on each organoid due to cell number limitations. First, we created a spatial network with $k_{\text{geom}} = 10$. We then identified spatially variable genes with the `binSpect` function, keeping the top 100 genes based on score. We then ran HMRF, setting the number of clusters to be identified to 10. The HMRF regularisation parameter ($\beta$) was for values ranging over $\beta \in \{0, 4, 8, \ldots, 48\}$ with 10 clusters for $\beta = 0, 4, \ldots, 48$ (13 settings), and we selected values from within a range where the domains were stable to variation in $beta$.

#### 2.5 Human DLPFC 10x Visium Data

The 10x Visium data of the dorsolateral prefrontal cortex (DLPFC) was obtained from the spatialLIBD project (`http://spatial.libd.org/spatialLIBD`). The data comprise 12 samples obtained from 3 subjects (4 samples per subject). The layers of each sample were manually annotated [8], with each sample having either 5 or 7 layers. We ran BANKSY on each sample separately, first normalizing the data to the median library size for each sample. Highly variable genes were then identified by modelling the mean-variance relationship for each gene as implemented in the Seurat package [11]. The data were subset to the top 2000 variable genes. Next, the neighborhood-augmented matrices were computed with a $1/r$ kernel, $\lambda = 0.15$ and $k_{\text{geom}} = 18$, corresponding to taking all spots up to second-order neighbours of a given index spot. After gene-wise z-scaling and dimensionality reduction of the BANKSY matrix, we performed Leiden clustering on the first 30 PCs and adjusted resolution and $k_{\text{expr}}$ parameters such that the number of clusters obtained matched the number of layers present in the manual annotation. Since more than one pair of resolution and $k_{\text{expr}}$ settings could yield the correct number of layers, for each dataset we report the median ARI across all parameter settings that had the correct layer number. Across all 12 datasets, we obtained a median ARI of 0.529.

For benchmarking, we ran Giotto's HMRF, BayesSpace and SpaGCN on the same data. For HMRFs, we used Giotto version 1.1.0, and followed the workflow on the package's site (`https://rubd.github.io/Giotto_site/articles/tut11_giotto_hmrf.html`). The workflow for all samples was the same. First, genes that were not detected in at least 10 cells were filtered out. Next the data was first count normalized to a fixed scale factor and log transformed, followed by gene- and cell-wise z-scaling. A spatial network was created with the Delaunay method and the maximum distance cutoff parameter was set to 'auto'. We identified genes with a spatially coherent expression pattern using the `binSpect` method with default parameters, and selected genes with adjusted $p$-value $< 0.1$ and with `binSpect` scores in the top 1 percentile, yielding approximately 160 genes per sample. HMRF was run on the spatial network and spatial genes identified, with the number of domains set to the number of layers present in the manual annotation. Based on the author's recommendations (`https://cran.r-project.org/web/packages/smfishHmrf/smfishHmrf.pdf`), we tested $\beta$ values (HMRF regularization parameter) from 0 to 100 in increments of 2. We then selected $\beta$ = 10, which gave the best median ARI across all samples (ARI = 0.380).

For BayesSpace (version 1.0.0), we followed the authors' vignette (`https://edward130603.github.io/BayesSpace/articles/maynard_DLPFC.html`). The workflow for all samples was the same. Highly variable genes were first identified by modelling the mean-variance relationship for each gene as implemented in the Scran package [7]. PCA was then performed on the top 2000 variable genes. We ran BayesSpace on the first 15 PCs, with the number of clusters set to the number of layers present in the manual annotation. The t-distributed error model was used, with 50,000 MCMC iterations, a burn-in period of 1000 iterations, and a gamma smoothing parameter of 3. Default values were used for all other parameters. We obtained a median ARI of 0.493 across all 12 samples.

For SpaGCN, we followed the authors' tutorial notebook (`https://github.com/jianhuupenn/SpaGCN/blob/master/tutorial/tutorial.ipynb`) up to Section 5.4 therein ('Plot spatial domains'), which showed analysis on sample 151673. We adapted the notebook for the 11 other samples, changing only the number of desired clusters but keeping all other parameters the same. While the clustering results obtained were not identical to what was reported in [4], comparisons of ARIs obtained showed close correspondence to reported values (Supp. Fig. 19c). We obtained a median ARI of 0.412 across all 12 samples.

### 3 Metagene Computation

Metagene expression captures the difference between two clusters that are to be compared, and can be used to visualise how BANKSY helps to better separate out the cells in subclusters in the neighbour-augmented space, relative to the original own-expression space (as shown in Figs. 2c and 3e, h). For the $i$-th cell, we define the metagene expression of sub-cluster 2 relative to subcluster 1 as the difference between the average expressions of DE genes more highly expressed in sub-cluster 2 and those more highly expressed in sub-cluster 1. Explicitly, for the $i$-th cell, if $\{g_{11}^{(i)}, \ldots, g_{1p}^{(i)}\}$ are the expression values (z-scaled over all the cells in the two sub-clusters) of each of the $p$ DE genes upregulated in subcluster 1, and similarly, $\{g_{21}^{(i)}, \ldots, g_{2q}^{(i)}\}$ are the values of the $q$ genes upregulated in sub-cluster 2, the metagene of the own-expression for sub-cluster 2 relative to sub-cluster 1 is defined as

$$m_{21}^{(i)} = \frac{\sum_{k=1}^{q} g_{2k}^{(i)}}{q} - \frac{\sum_{j=1}^{p} g_{1j}^{(i)}}{p}. \tag{3}$$

The metagene expression for the neighbour expression is defined analogously using the corresponding $p$ and $q$ neighbour expression rows of these genes in the neighbour-augmented matrix. Once the metagene expressions are computed, each cell can be plotted in the the product space of the neighbour versus own-expression metagene.

### 4 Cluster Consensus Across Runs

In several analyses, we had to match the cluster labels across different clustering solutions. We did this by posing the cluster matching problem as a linear sum assignment problem (also known as the weighted bipartite matching problem), which admits a strongly polynomial solution via the Hungarian algorithm [5]. Briefly, given two clusterings, we set up a matrix such that the rows and columns corresponded to the clusters in the two clusterings respectively, and the entries were the number of cells that were common between the pair of clusters. The goal was to find, for each row of this matrix, a unique matching column, so that the number of cells that were common between the matching clusters was maximised over all the clusters. For the Python version of the code, we used the Scipy implementation (`scipy.optimize.linear_sum_assignment`) to match clusters across different clustering runs. For the R version of the code, we used the `HungarianSolver` function within the `RcppHungarian` library on CRAN.

All in all, this matching allowed us to visually compare different parameter settings and match clusters from BANKSY clustering to their closest equivalents in the non-spatial case (or other methods that we compared to, like BayesSpace, SpaGCN, HMRF, etc.). For example, the matching of the clusters in Fig. 4 between the non-spatial clustering, BANKSY clustering and MERINGUE clustering was achieved using this methodology.

### References


[1] *Allen Mouse Brain Atlas [dataset]*. 2011. URL: `http://mouse.brain-map.org`.

[2] Dylan M. Cable et al. "Robust decomposition of cell type mixtures in spatial transcriptomics". In: *Nature Biotechnology* (Feb. 2021). ISSN: 1546-1696. DOI: `10.1038/s41587-021-00830-w`.

[3] Marie Chavent, Vanessa Kuentz-Simonet, Amaury Labenne, and Jérôme Saracco. "ClustGeo: an R package for hierarchical clustering with spatial constraints". In: *Computational Statistics* 33.4 (Dec. 2018). arXiv: 1707.03897, pp. 1799–1822. ISSN: 0943-4062, 1613-9658. DOI: `10.1007/s00180-018-0791-1`.

[4] Jian Hu et al. “SpaGCN: Integrating gene expression, spatial location and histology to identify spatial domains and spatially variable genes by graph convolutional network”. In: *Nature Methods* 18.11 (Nov. 2021), pp. 1342–1351. ISSN: 1548-7105. DOI: 10.1038/s41592-021-01255-8.

[5] H. W. Kuhn. “The Hungarian method for the assignment problem”. en. In: *Naval Research Logistics Quarterly* 2.1-2 (1955), pp. 83–97. ISSN: 1931-9193. DOI: 10.1002/nav.3800020109.

[6] Ed S. Lein et al. “Genome-wide atlas of gene expression in the adult mouse brain”. en. In: *Nature* 445.7124 (Jan. 2007), pp. 168–176. ISSN: 1476-4687. DOI: 10.1038/nature05453. (Visited on 04/11/2022).

[7] Aaron T. L. Lun, Davis J. McCarthy, and John C. Marioni. *A step-by-step workflow for low-level analysis of single-cell RNA-seq data with Bioconductor*. en. Tech. rep. 5:2122. Type: article. F1000Research. DOI: 10.12688/f1000research.9501.2.

[8] Kristen R. Maynard et al. “Transcriptome-scale spatial gene expression in the human dorsolateral prefrontal cortex”. In: *Nature Neuroscience* 24.3 (Mar. 2021), pp. 425–436.

[9] Brendan F Miller et al. “Characterizing spatial gene expression heterogeneity in spatially resolved single-cell transcriptomics data with nonuniform cellular densities”. In: *Genome Research* (2021). DOI: 10.1101/gr.271288.120.

[10] Jeffrey R. Moffitt et al. “Molecular, spatial, and functional single-cell profiling of the hypothalamic preoptic region”. en. In: *Science* 362.6416 (Nov. 2018). DOI: 10.1126/science.aau5324.

[11] Tim Stuart et al. “Comprehensive Integration of Single-Cell Data”. In: *Cell* 177.7 (June 2019), 1888–1902.e21. ISSN: 0092-8674. DOI: 10.1016/j.cell.2019.05.031.

[12] V. A. Traag, L. Waltman, and N. J. van Eck. “From Louvain to Leiden: guaranteeing well-connected communities”. In: *Scientific Reports* 9.1 (Mar. 2019), p. 5233. ISSN: 2045-2322. DOI: 10.1038/s41598-019-41695-z.

[13] *Vizgen Data Release Program*. May 2021. URL: https://vizgen.com/support/data-release-program.

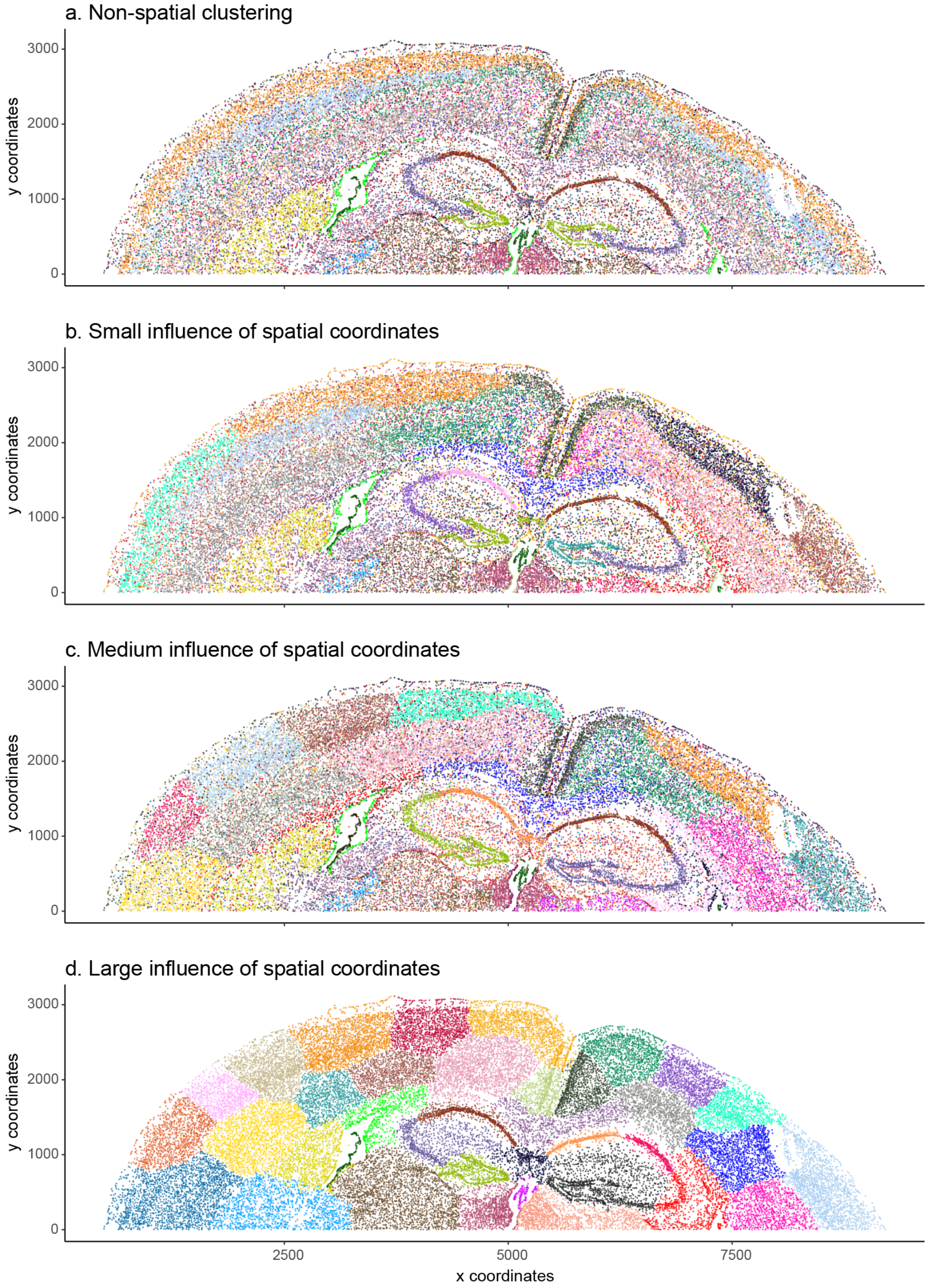


Figure 1: Clustering results with the spatial coordinates of cells appended to the gene expression vectors as features (i.e., using the incorrect approach to incorporating spatial information; Results section and Supp. Section 1). Coronal section of the mouse brain [13]. (a) Non-spatial clustering. As expected, the left and the right halves of the brain are similar, and are marked by the same clusters. (b) When the spatial locations of cells are appended to the cells' features, cells that are far apart in physical space are also far apart in feature space, even if they have the same transcriptomic signatures. Thus, the clustering algorithm labels them differently. (c, d) This effect is stronger when the weighting ($\lambda$) of the spatial coordinates as features is increased.

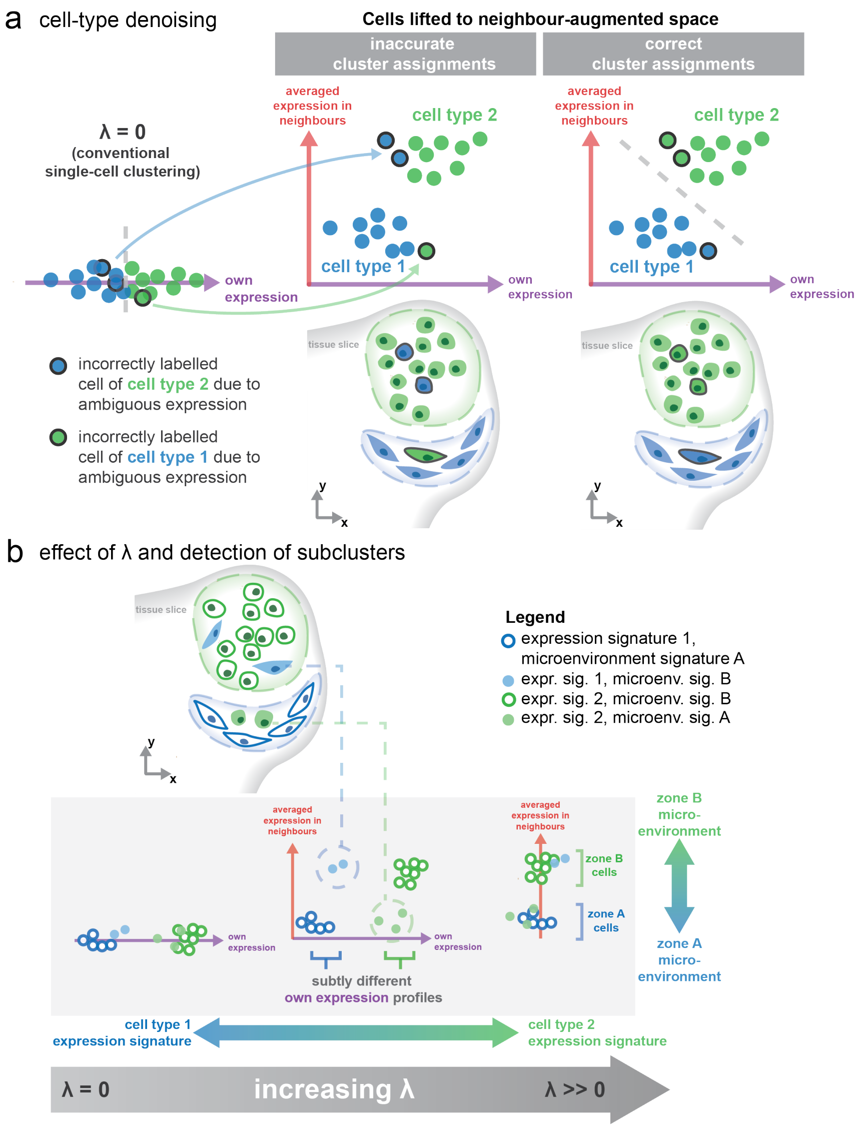


Figure 2: Detailed schematic to aid conceptual understanding of BANKSY. (a) How BANKSY may lead to more accurate cluster assignments relative to non-spatial clustering ($\lambda = 0$) by accounting for local miroenvironmental information. Cells near the decision boundary in own expression space alone get better separated in the neighbour-augmented space. (b) Top–Schematic of tissue slice shows physical locations of two cell types (1 and 2) in two zones (A and B). Zone A contains mainly cell type 1 but some of cell type 2. Zone B contains mainly cell type 2 but some of cell type A. Bottom–Effect of increasing $\lambda$: at $\lambda = 0$ (bottom left), only the cell's transcriptome is used, making it identical to conventional unsupervised clustering analysis. As $\lambda$ increases (bottom middle), subsets of cells in different environments with subtly different transcriptomes become easier to tease out. For instance, the blue circles–filled and empty–correspond to the same cell type in two different neighbourhoods, possessing subtle transcriptomic differences. A similar effect is also shown for the green cell type. At higher lambdas (bottom right), a zone or spatial-domain finding effect occurs where zones representing different microenvironments are clustered separately, and can comprise multiple cell types per zone.

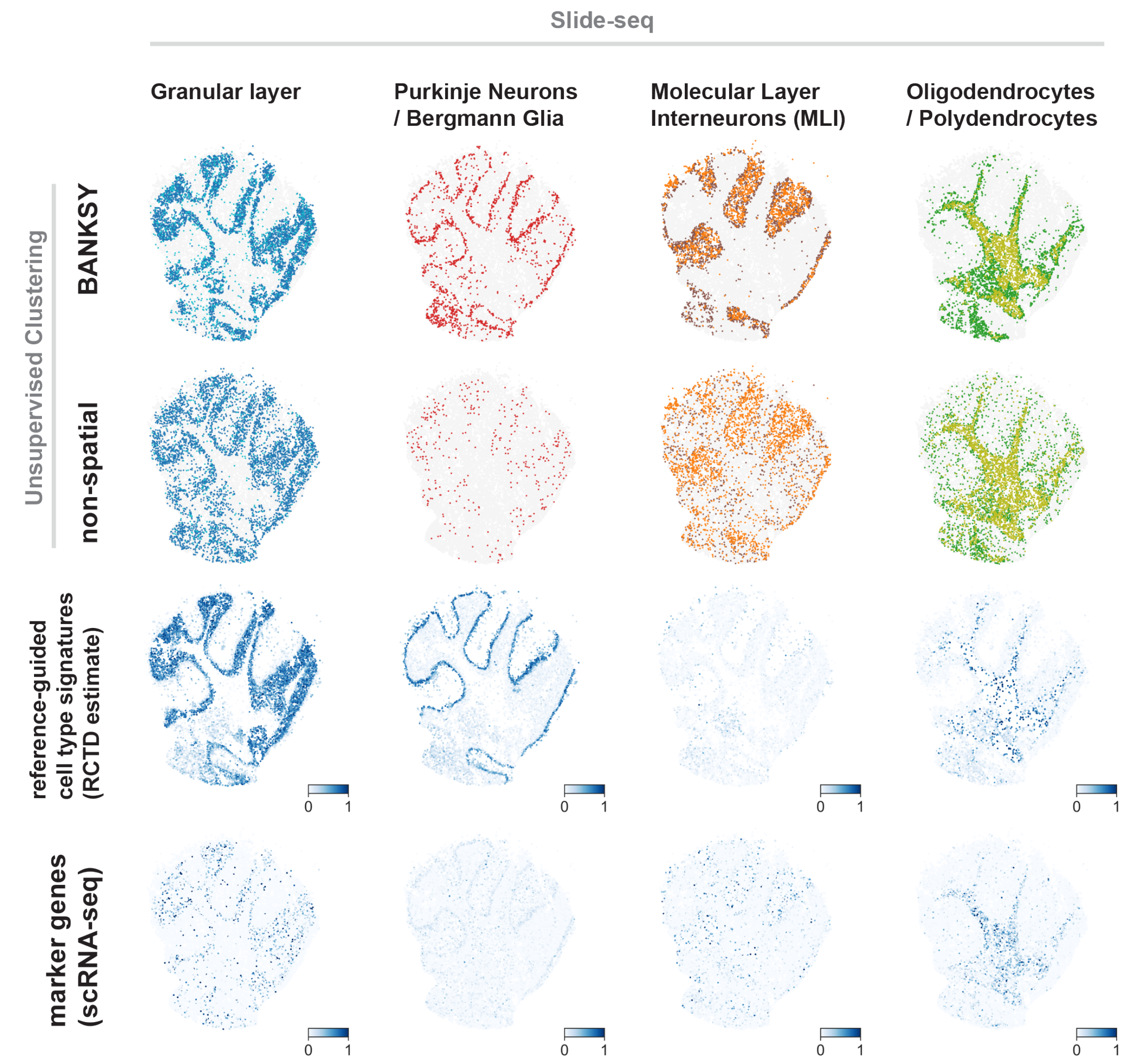


Figure 3: Cluster assignments for major clusters in Slide-seq cerebellum dataset. Granular layer for BANKSY consists of two sub-clusters, coloured in blue and cyan. For non-spatial clustering, no cluster matched the second cluster in BANKSY. Molecular layer interneurons (MLI) consist of two sub-clusters, coloured in orange and brown. Oligodendrocytes/polydendrocytes consist of two sub-clusters, coloured in green and olive. Lower two rows show comparison to RCTD weights from corresponding clusters in scRNA-seq reference dataset, and top DE marker genes from corresponding clusters in the reference dataset (obtained from dropviz.org).

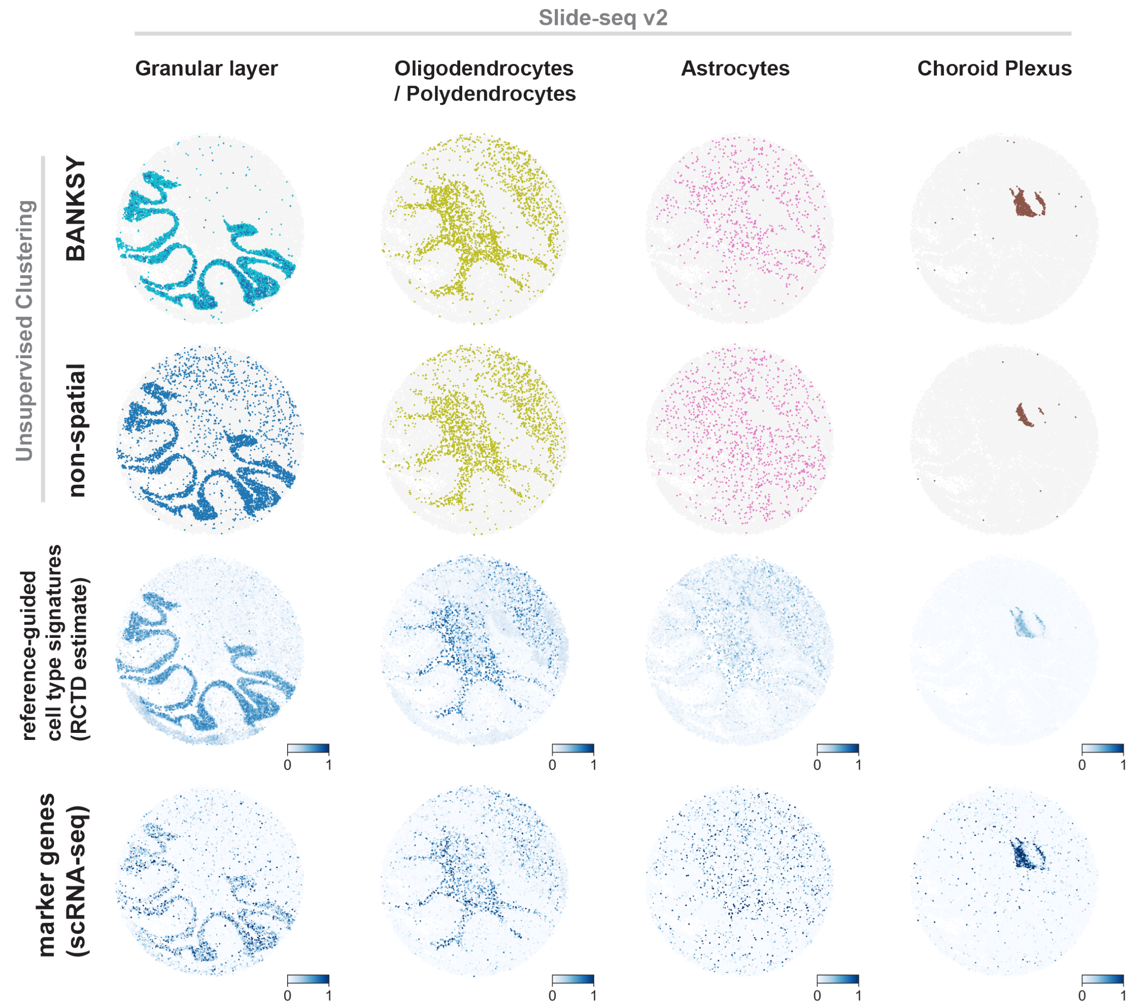


Figure 4: Cluster assignments for major clusters including oligodendrocytes/polydendrocytes, astrocytes and choroid plexus not shown in Fig. 2. Granular layer consists of two sub-clusters, coloured blue and cyan. Lower two rows show comparison to RCTD weights from corresponding clusters in scRNA-seq reference dataset, and top DE marker genes from corresponding clusters in the reference dataset (obtained from dropviz.org).

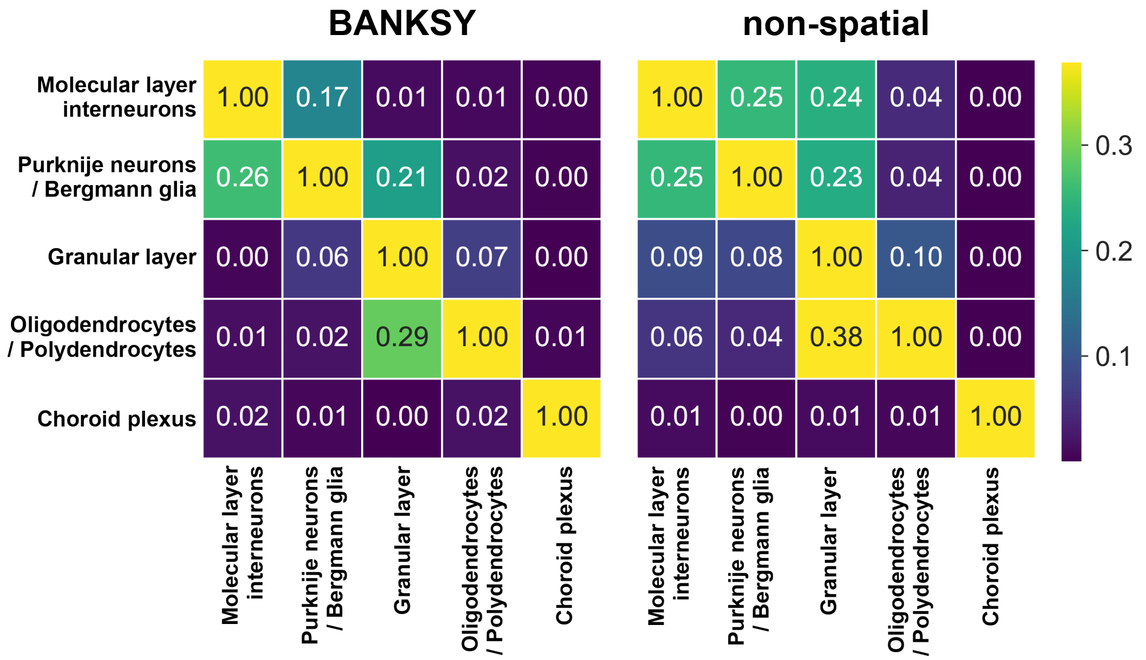


Figure 5: Normalised connection scores across layers for non-spatial and BANKSY clustering in Slide-seq v2 cerebellum dataset. Higher scores indicate greater intermingling between layers. Colourmap is clipped to highest cross-layer value (0.38).

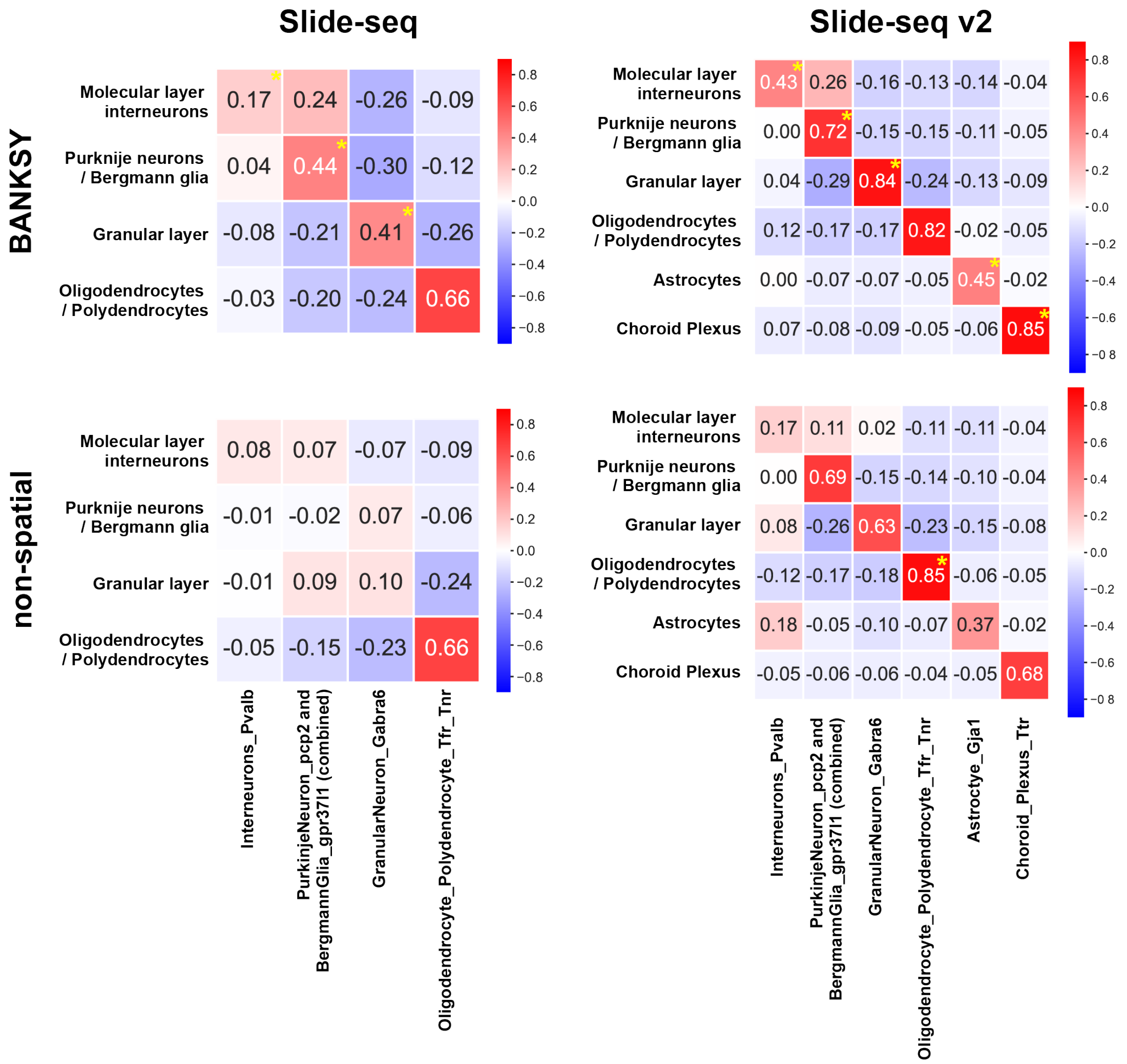


Figure 6: Point-biserial correlation ($r$) between each major cluster and RCTD weights from corresponding reference cluster when using BANKSY clustering (a) and using unsupervised clustering without spatial information (b), for both Slide-seq and Slide-seq v2. Asterisks along diagonals indicate higher $r$ values. BANKSY yields higher correlations for all major clusters in both datasets except for the oligodendrocyte/polydendrocyte cluster. All correlations along the diagonals are significant ($p < 0.05$).

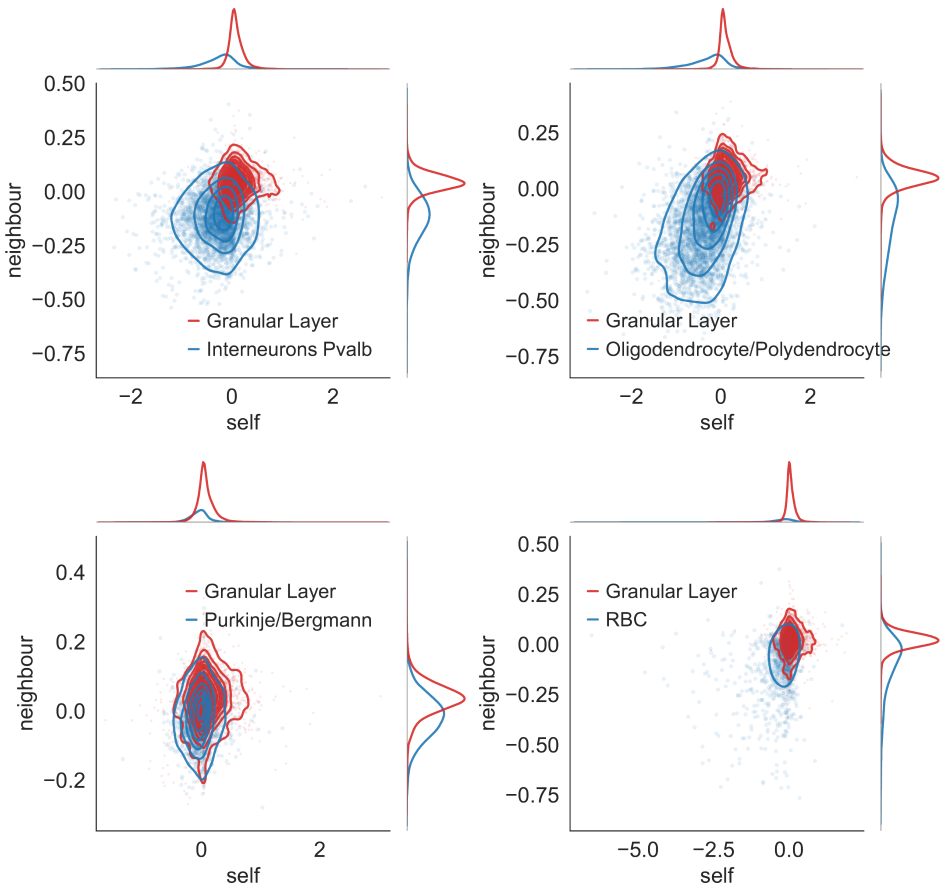


Figure 7: Plot of difference in metagenes (mean of top 20 DE genes, see Supp. Section 3) in the granular layer cluster vs other clusters in Slide-seq, comparing each other cluster (blue) to the granular layer (red). The additional information provided by neighbour expressions leads to clearer separation in own expression-neighbour expression metagene space.

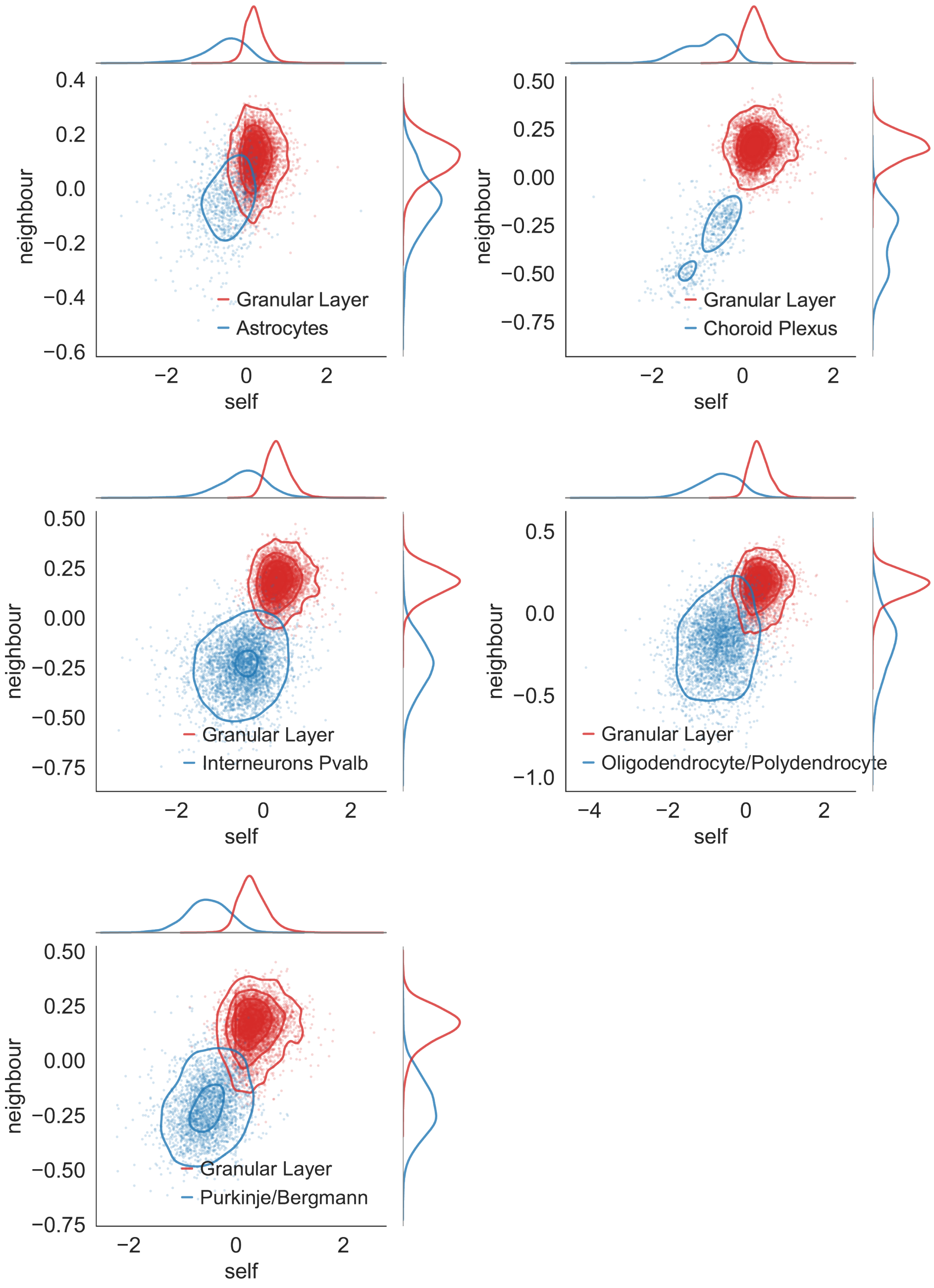


Figure 8: Plot of difference in metagenes (mean of top 20 DE genes, see Supp. Section 3) in the granular layer cluster vs other clusters in Slide-seq v2, comparing each other cluster (blue) to the granular layer (red). The additional information provided by neighbour expressions leads to clearer separation in own expression-neighbour expression metagene space.

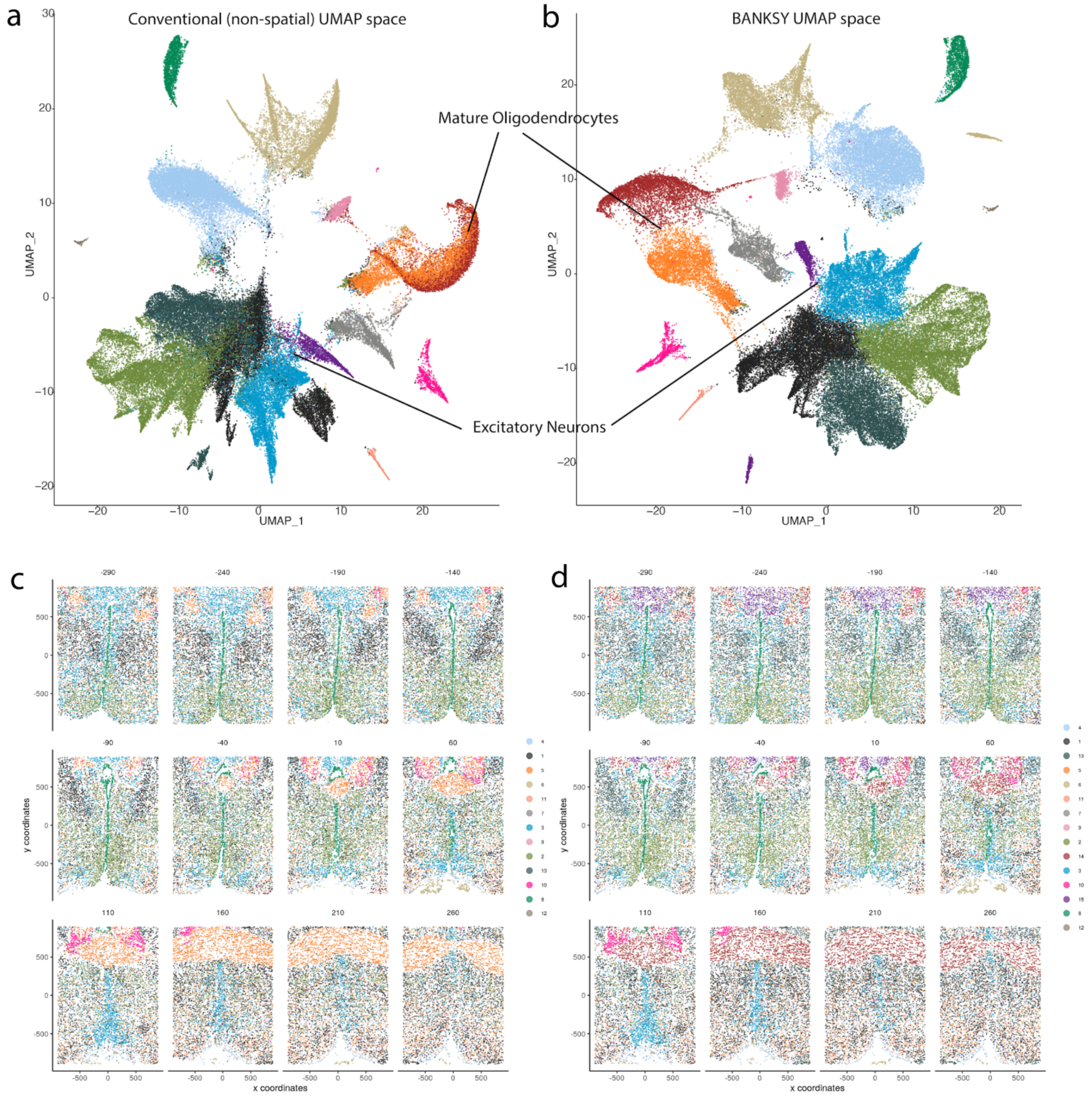


Figure 9: (a) UMAP representation of the cells in the own expression feature space (which is all that is used for non-spatial clustering). (b) UMAP representation of cells in the BANKSY (neighbour-augmented) feature space. Both UMAPs (a) and (b) are coloured by the BANKSY clustering labels, and show that the mature oligodendrocyte sub-clusters are mixed in the own-expression (conventional) feature space, but separate out well in the neighbour-augmented space (orange and red cluster). This effect is less pronounced for the excitatory neurons (blue and purple clusters). (c, d) The tissue maps show all twelve z-planes with cells colored by both non-spatial clustering (c) and BANKSY clustering (d).

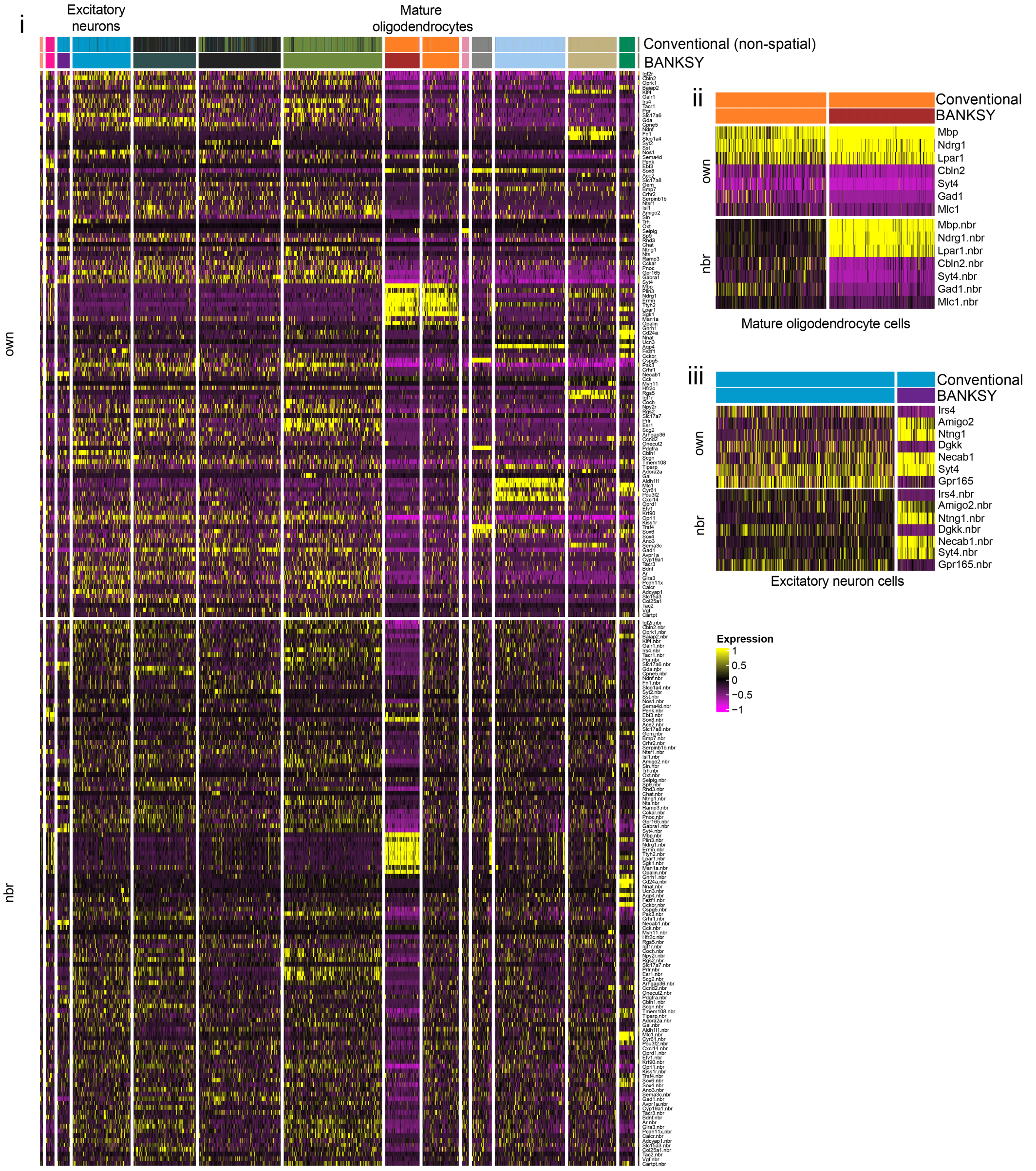


Figure 10: Heatmaps showing expression of genes in the mouse hypothalamus data shown in Figure 3. (i) Heatmap showing all 73655 cells, with cluster labels given for conventional (non-spatial) clustering and BANKSY clustering ($\lambda = 0.2$) in cell-typing mode. The expression levels shown were z-scaled across all 73655 cells. (ii) Subset of cells and genes from the heatmap in (i), but only showing the cells labelled as mature oligodendrocytes in both conventional clustering and BANKSY, and showing the top DE genes between the two BANKSY sub-clusters (orange and red, corresponding to sub-clusters 1 and 2 in Fig. 3c). The heatmap expression levels are the same as those shown in the full heatmap in (i), and show that the difference in cells' own expressions between the two sub-clusters is small, but that in their neighbourhoods (bottom half of heatmap) is large. This explains why conventional clustering is not able to identify these sub-clusters based only on the differences in their own expression, but BANKSY clustering is able to resolve these differences. (iii) Similar to the heatmap in (ii), but for the excitatory neurons. The differences in own expression are greater here, but utilising neighbourhood information still helps to separate them more easily.

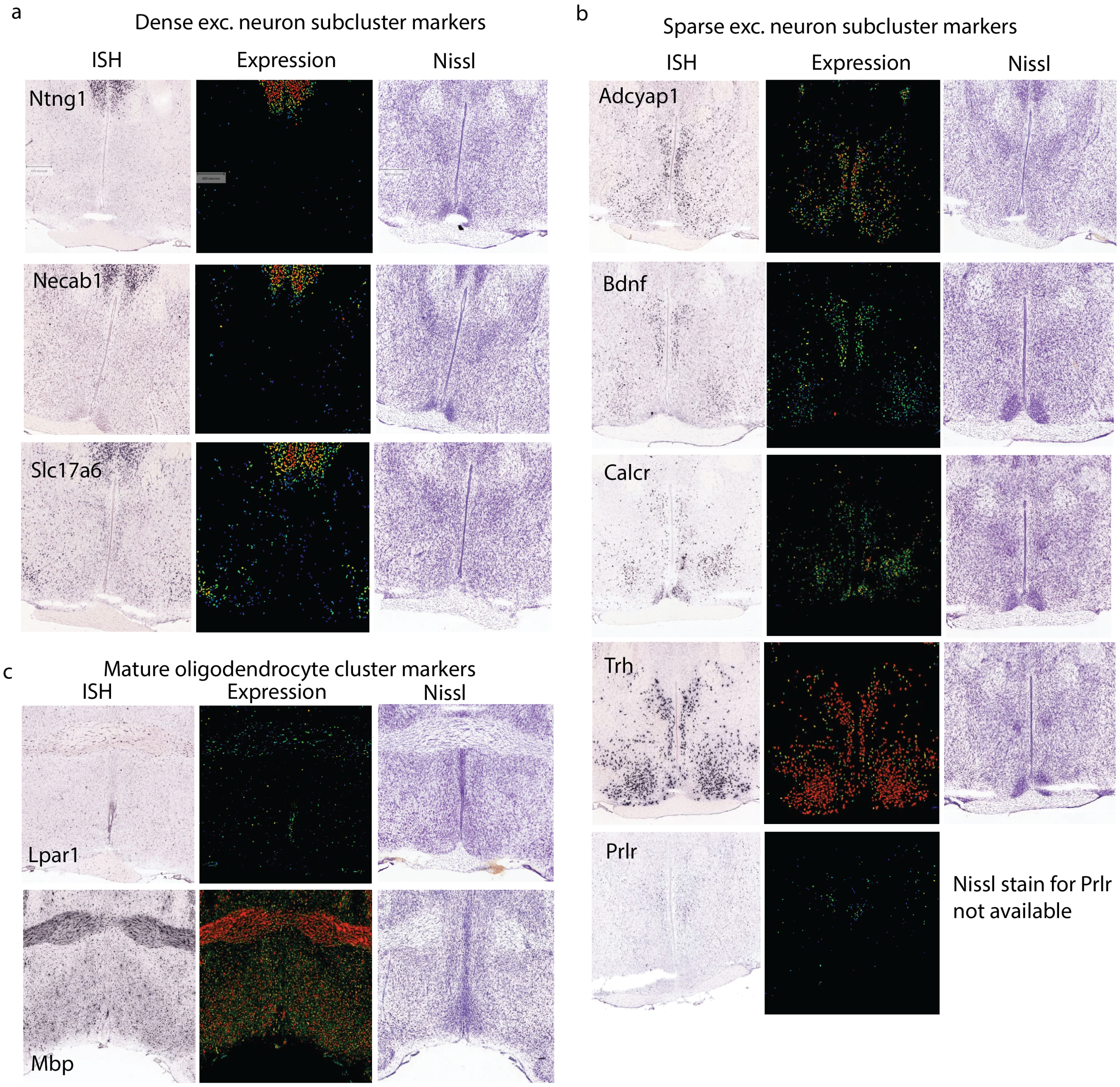


Figure 11: (a) Allen Mouse Brain Atlas [1, 6] images showing some of the genes that are differently expressed within the mature oligodendrocyte sub-clusters and the excitatory neuron sub-clusters in Fig. 3.

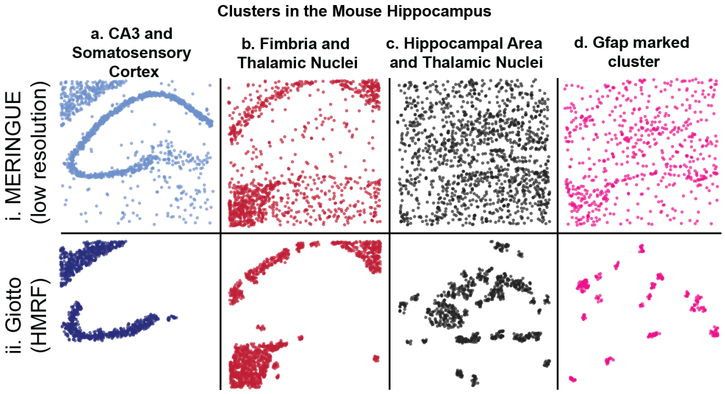


Figure 12: MERINGUE at low resolution and HMRF applied to the mouse hippocampus data (Fig. 4). MERINGUE (low resolution) was computed with $k_{\text{expr}} = 50$ (the number of neighbours in the graph in expression space), while the value of $k_{\text{expr}}$ for the MERINGUE results shown in Fig. 4 was 15 (see Supp. Section 2.2 for details about the two MERINGUE runs). (ii) Spatial regularisation in HMRF based methods like Giotto tends to encourage physically adjacent cells to be identically labelled, and are as such better suited for zone finding. Thus Giotto tended to perform poorly for cell typing, especially when a cell type was not tightly packed in physical space (c, d, ii) or multiple cell types are intermingled in the same tissue (as they were in the fimbria, the thalamic nuclei, and the general hippocampal area).

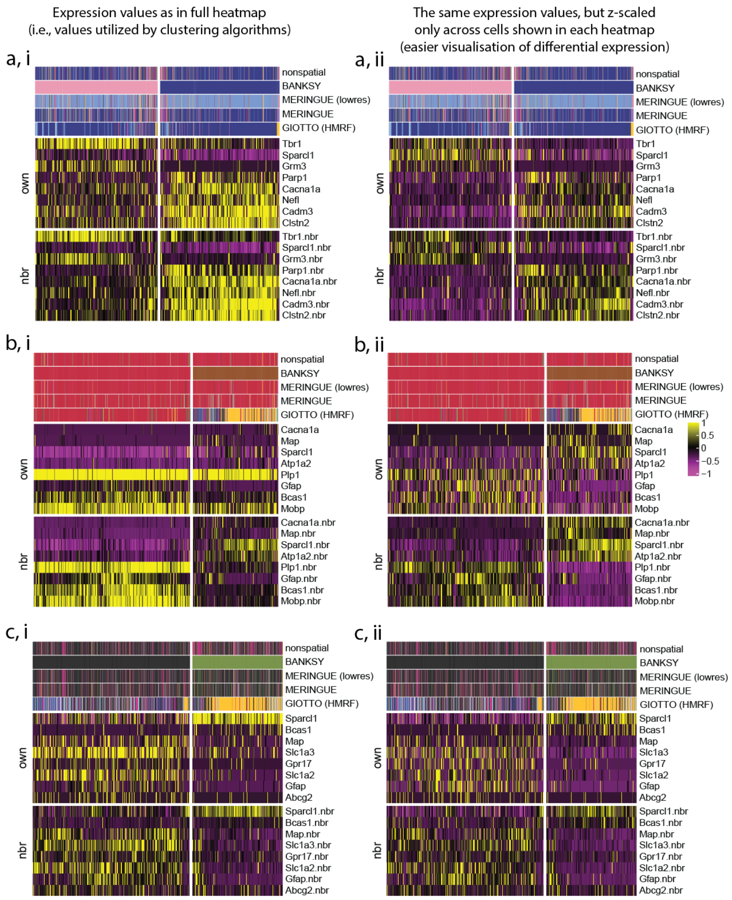


Figure 13: Heatmaps showing the expression of DE genes for the clusters distinguished by BANKSY in the mouse hippocampus data (Fig. 4). Cells are ordered by the BANKSY labels. The labels for non-spatial clustering, MERINGUE clustering (most closely matched to BANKSY and non-spatial clustering), MERINGUE clustering with parameters matched to the non-spatial and BANKSY clustering (which has a lower effective clustering 'resolution' than these, Supp. Section 2.2), and Giotto's HMRF labelling are also shown with annotation bars. (a, i) Genes that are DE between the somatosensory cortex cluster and CA3 cluster labelled by BANKSY. Expression levels are not z-scaled across just these cells, and are instead z-scaled across all cells in this dataset (i.e., the set of cells shown in the heatmap in Supp. Fig. 16a). To better visualise the differences in the expression of these genes between just these two clusters, we z-scaled the expression of each gene in just the cells in these two clusters, as shown in (a, ii). (b) Same as (a), but for the fimbria and thalamic nuclei clusters shown in Fig. 4b. (c) Same as (a), but for the general hippocampal area and thalamic nuclei clusters shown in Fig. 4c.

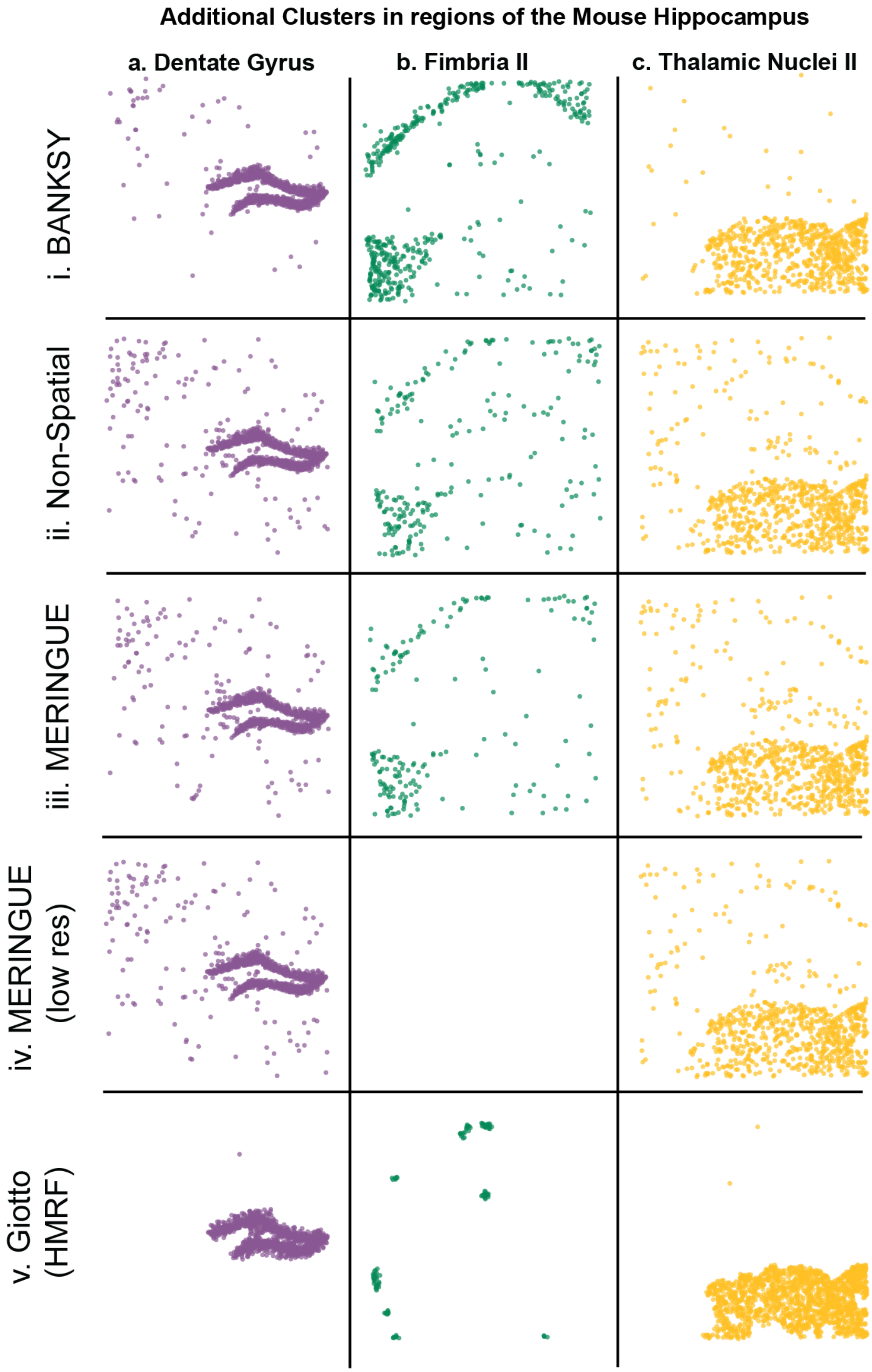


Figure 14: Additional clusters from the mouse hippocampus data shown in Fig. 4, comparing the different clustering methods.

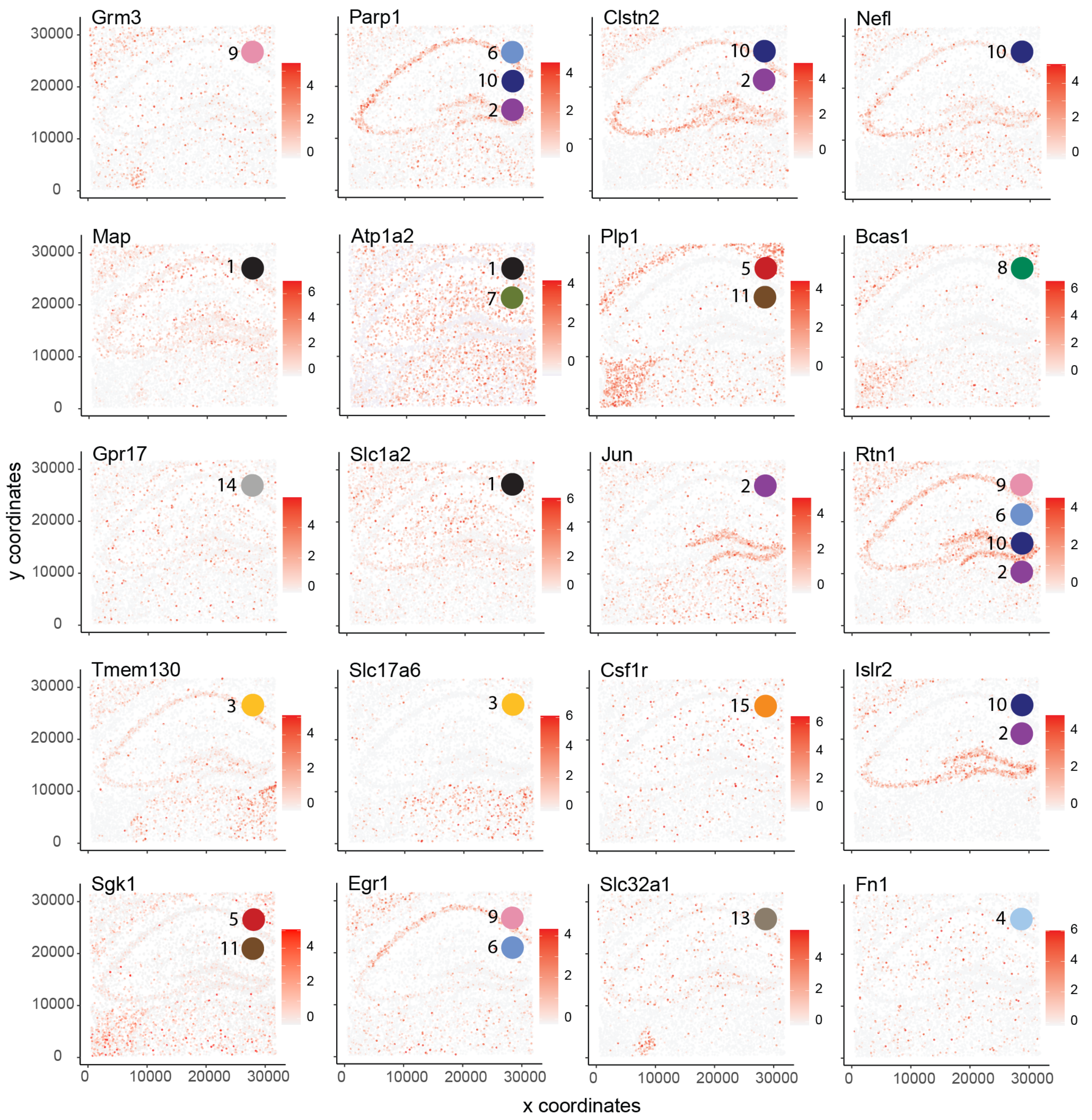


Figure 15: Spatial distribution of genes that are DE between the pairs of BANKSY clusters found in the mouse hippocampus data in Fig. 4. Red: high expression, White: low expression. Coloured circles mark the cluster colours that the corresponding gene is more highly expressed in, relative to the other cluster in that pair. Numbers next to the coloured circles represent the cluster ID. For instance, Grm3 is more highly expressed in the somatosensory cortex cell cluster (light pink) in Fig. 4a, i, compared to the CA3 pyramidal cell cluster (dark blue) and therefore distinguishes these two clusters. For clusters that are not part of a pair, such as the dentate gyrus cluster (dark purple in Supp. Fig. 14), a circle means that that gene (such as Parp1, or Clstn2) is highly expressed in that cluster.

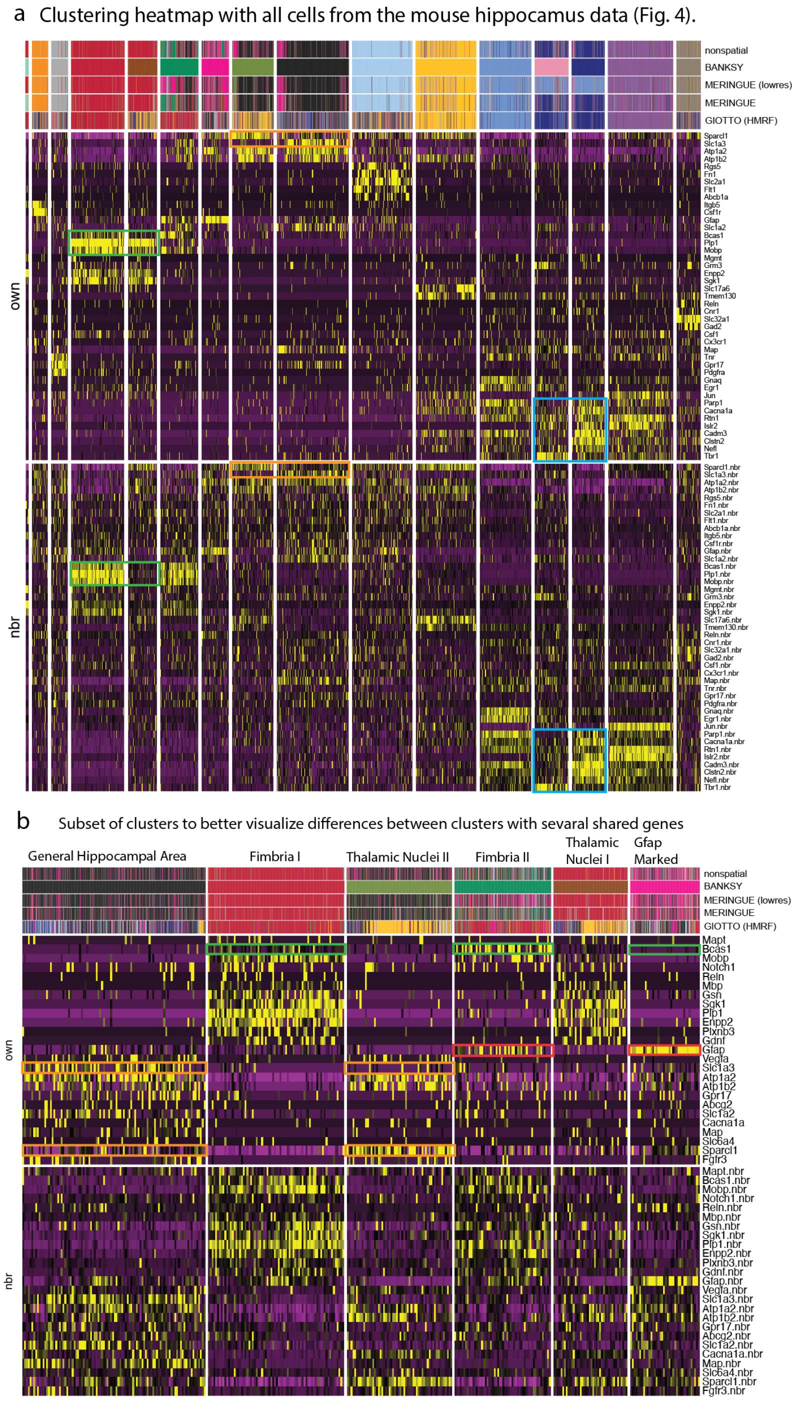


Figure 16: Heatmaps showing the own and neighbour expression for the mouse hippocampus data in Fig. 4. (a) Heatmap with all cells in the dataset. Boxes show the own and neighbourhood expressions of some of the genes that are DE between the clusters distinguished by BANKSY in Fig. 4. (b) Heatmap showing a related subset of clusters. Orange boxes show that Slc1a3 and Sparcl1 distinguish the general hippocampal area cluster (black) and the thalamic nuclei cluster (olive green). Red boxes show the difference in the expression of Gfap between the green and pink clusters, and green boxes show that Bcas1 is strongly DE between these two clusters. Bcas1 (green boxes) also distinguishes the red and green clusters, which are both located in the fimbria region.

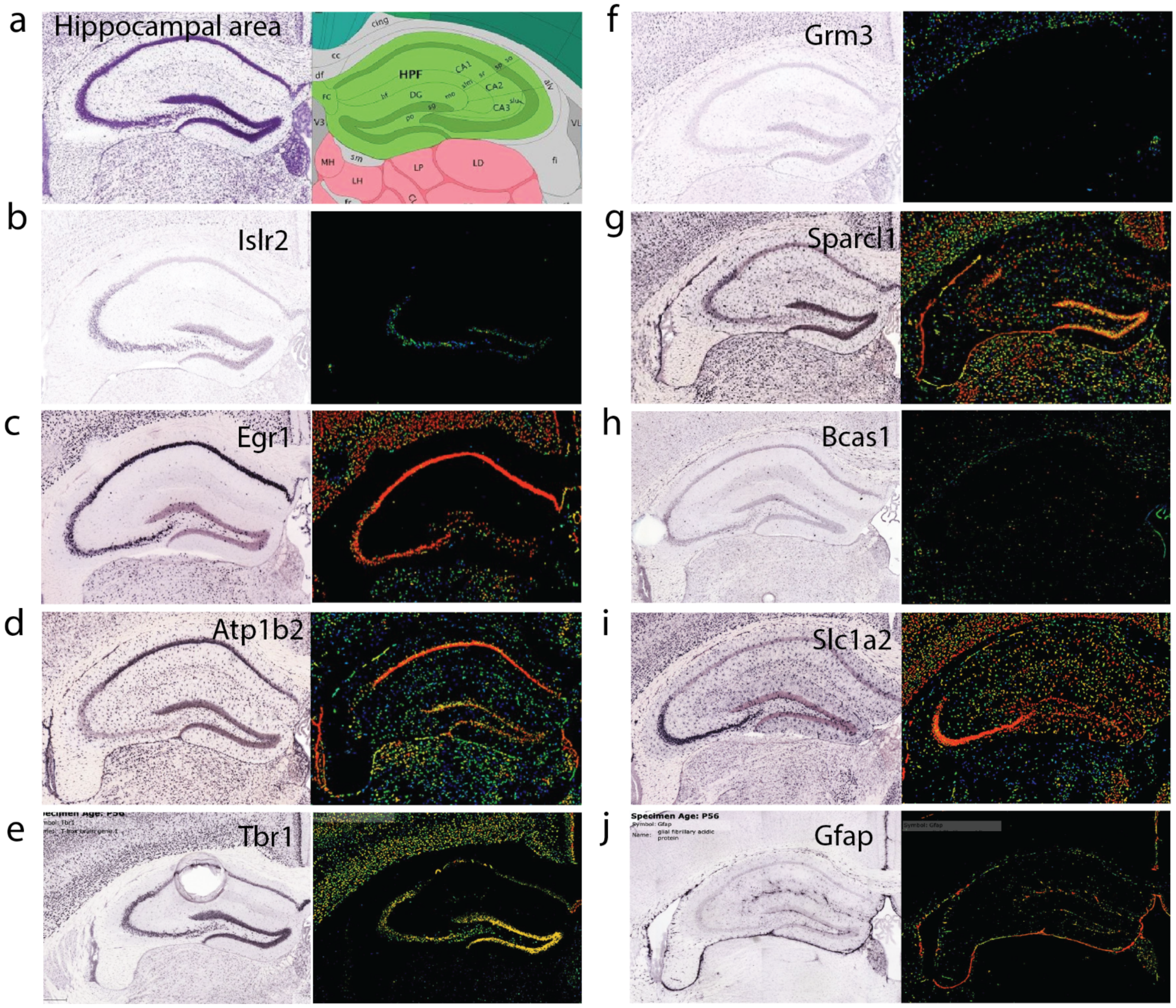


Figure 17: Allen Mouse Brain Atlas images [1, 6] showing ISH and expression levels for some of the DE genes in Fig. 4. (a) Schematic labelling the main regions in the hippocampus. (b-j) ISH images and expression levels for genes in this region.

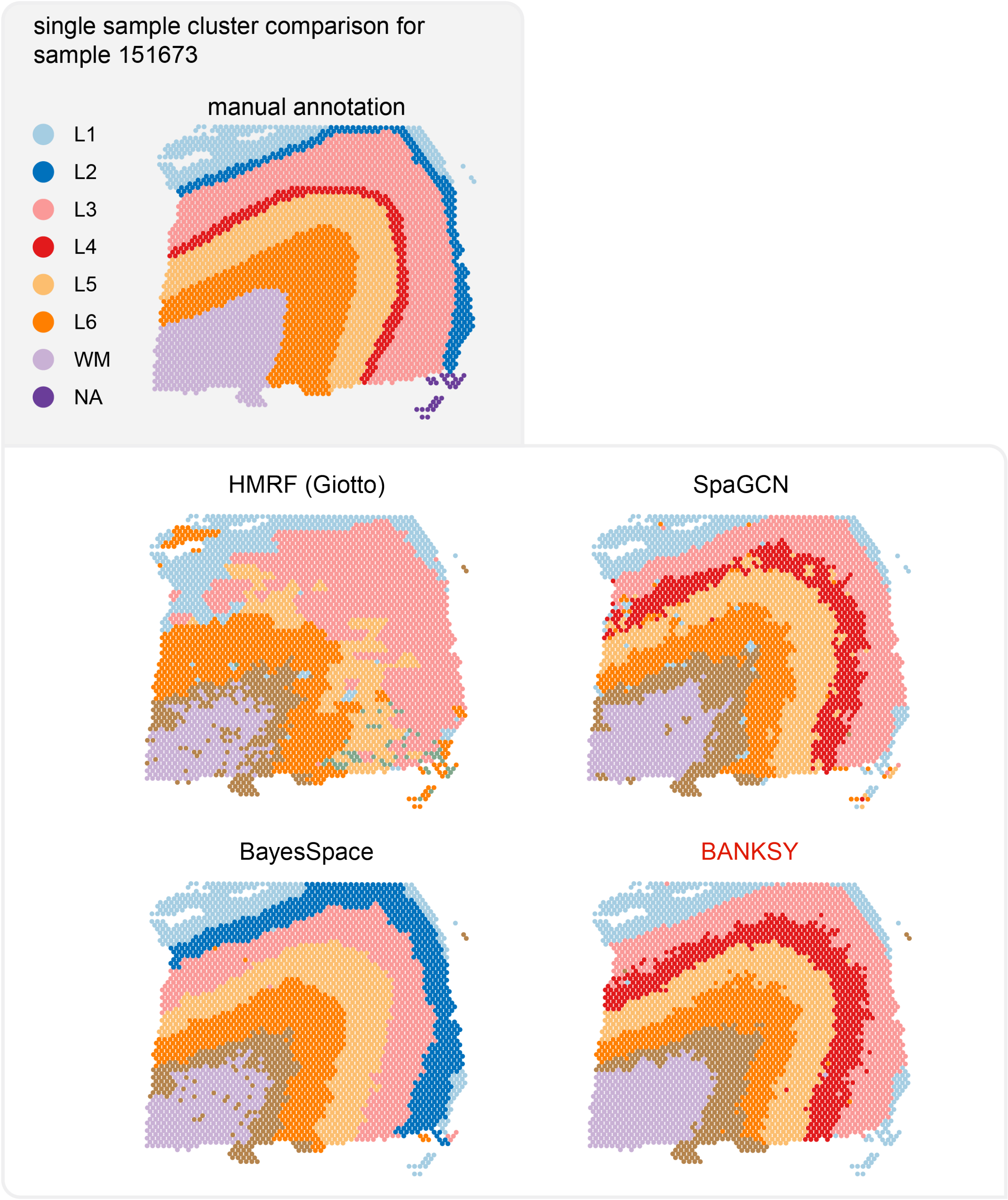


Figure 18: Qualitative cluster comparison for sample 151673.

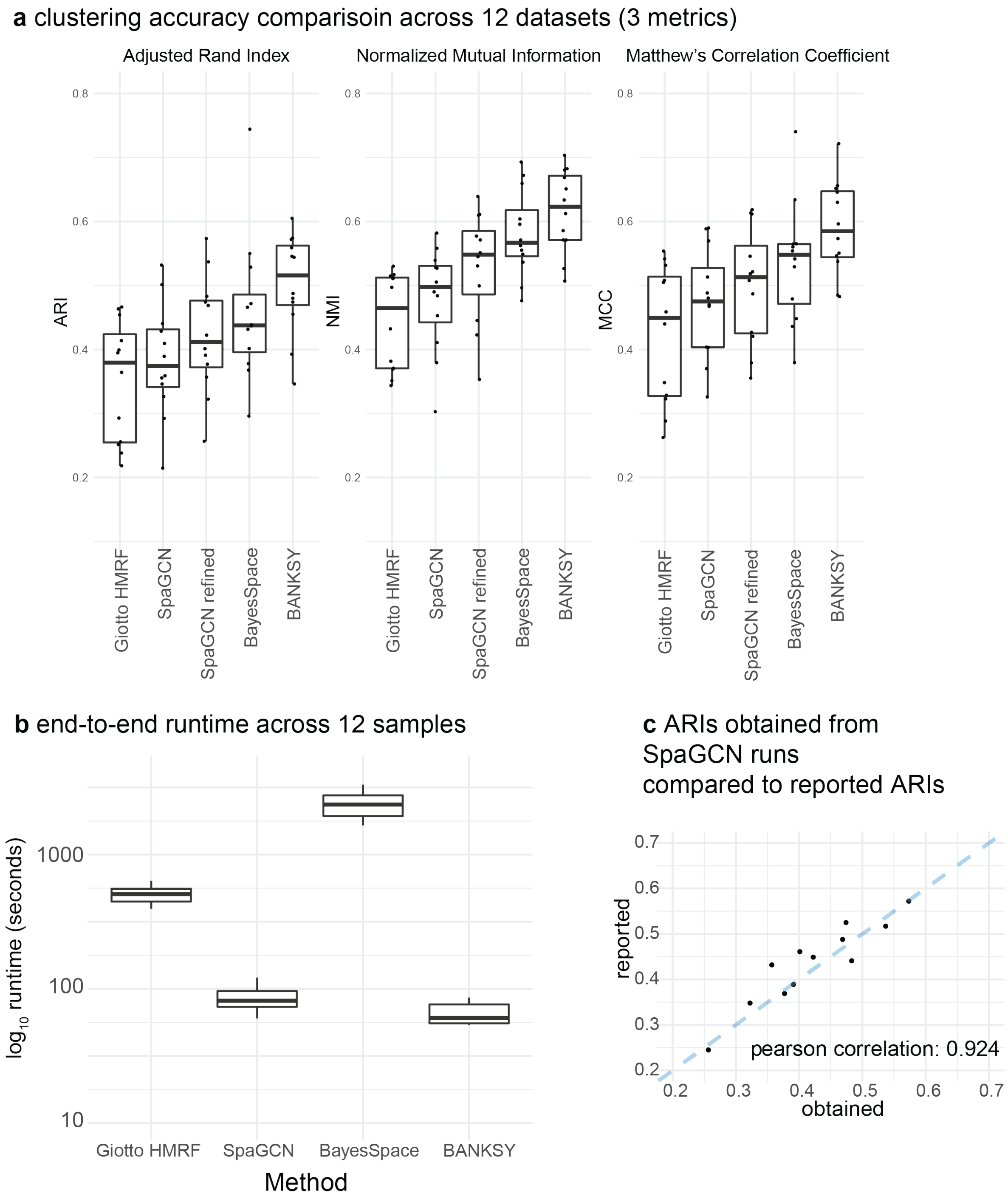


Figure 19: Supplementary figures for domain-finding comparison on 10X Visium Human DLPFC dataset. (a) Clustering accuracy comparisons using 3 different metrics: adjusted Rand index (ARI), normalized mutual information (NMI) and the Matthews correlation coefficient (MCC). SpaGCN and SpaGCN-refined show the SpaGCN method before and after the refinement step. Note that in Figure 6, SpaGCN after refinement was shown. (b) End to end run times for all 12 datasets with each method tested. (c) Results for our runs with SpaGCN using the example notebook provided with default parameters. Clusters were not identical but ARIs closely matched their reported values.

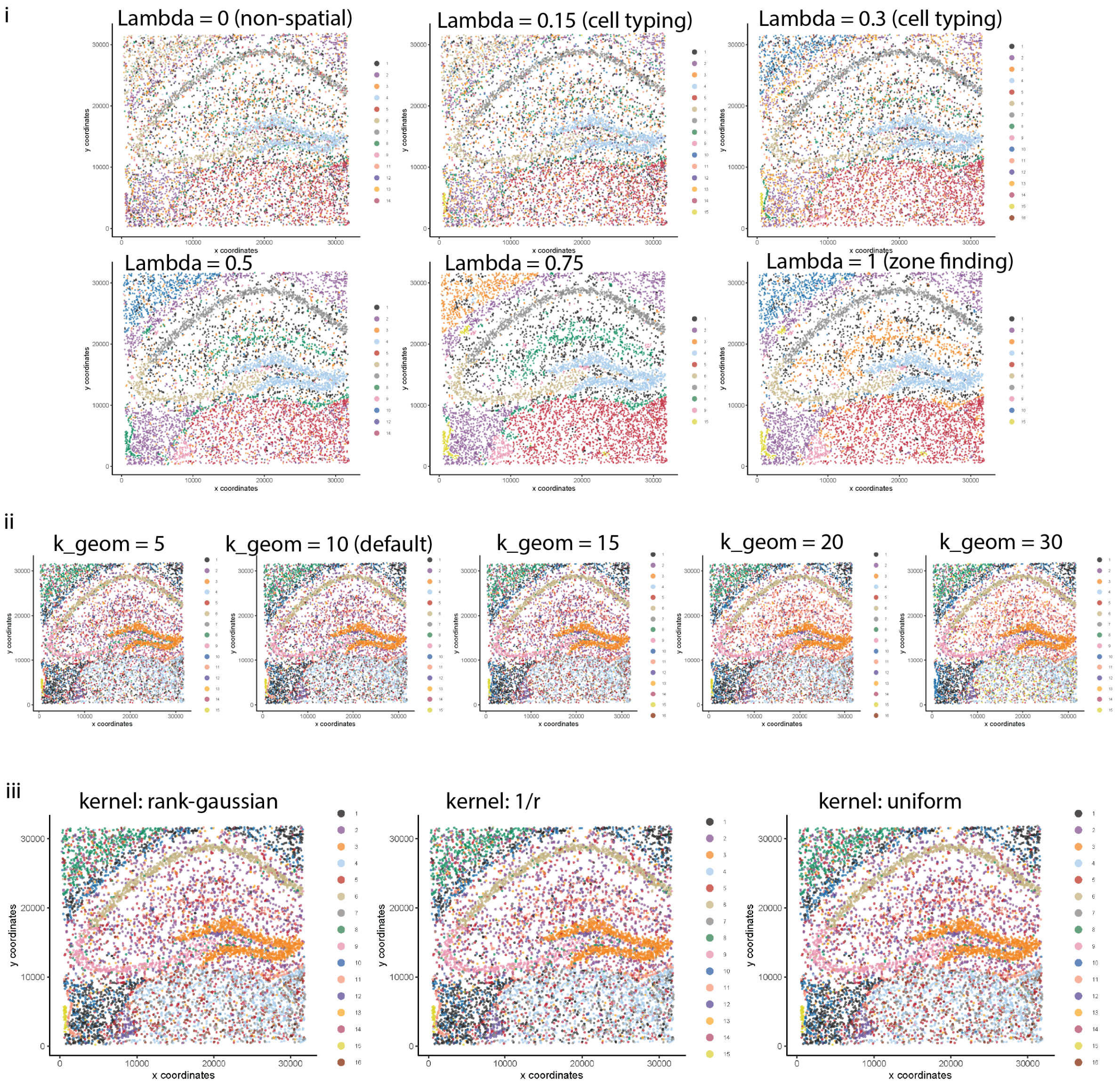


Figure 20: Parameter sweeps over the three major spatial parameters: (i) the mixing parameter $\lambda$, (ii) the number of spatial neighbours to average over to get neighbourhood signature ($k_{\text{geom}}$), and (iii) the weighting kernel to use when weighting spatial neighbours (rank-Gaussian, defined in Section 4.1, refers to a Gaussian kernel, but with the rank of a neighbour used instead of the distance, where, for instance, rank = 2 means it is the second closest neighbour to the index cell.)

i

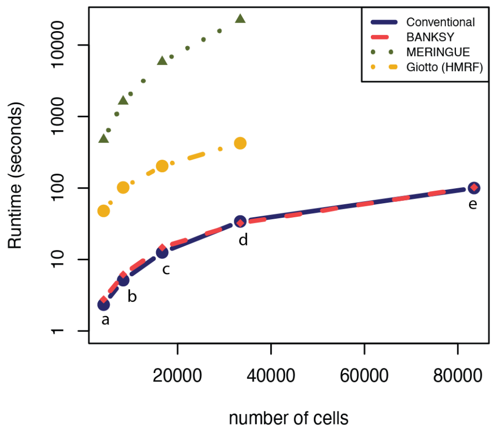


ii

| Num. Cells | Conventional (s) | BANKSY (s) | MERINGUE (s) | Giotto (s) |
|---|---|---|---|---|
| 4178 | 2.3 | 2.8 | 475.7 | 47.9 |
| 8356 | 5.1 | 6.2 | 1628.8 | 101.8 |
| 16709 | 12.6 | 14.8 | 5833.0 | 203.2 |
| 33419 | 34.1 | 32.3 | 22645.0 | 424.0 |
| 83463 | 100.0 | 102.3 | NA | NA |

| Num. cells | BANKSY matrix (s) | BANKSY clust (s) |
|---|---|---|
| 4178 | 0.4 | 2.4 |
| 8356 | 1.1 | 5.0 |
| 16709 | 2.0 | 12.8 |
| 33419 | 2.6 | 29.7 |
| 83463 | 8.3 | 94.0 |

| Num. cells | MERINGUE graph (s) | MERINGUE clust (s) |
|---|---|---|
| 4178 | 5.9 | 469.8 |
| 8356 | 14.4 | 1614.5 |
| 16709 | 32.0 | 5801.0 |
| 33419 | 71.4 | 22573.6 |

| Num. cells | Giotto network (s) | Giotto spat. genes (s) | Giotto E-M (s) |
|---|---|---|---|
| 4178 | 0.9 | 20.8 | 26.1 |
| 8356 | 2.5 | 42.3 | 57.0 |
| 16709 | 7.9 | 86.6 | 108.6 |
| 33419 | 27.2 | 177.4 | 219.3 |

iii

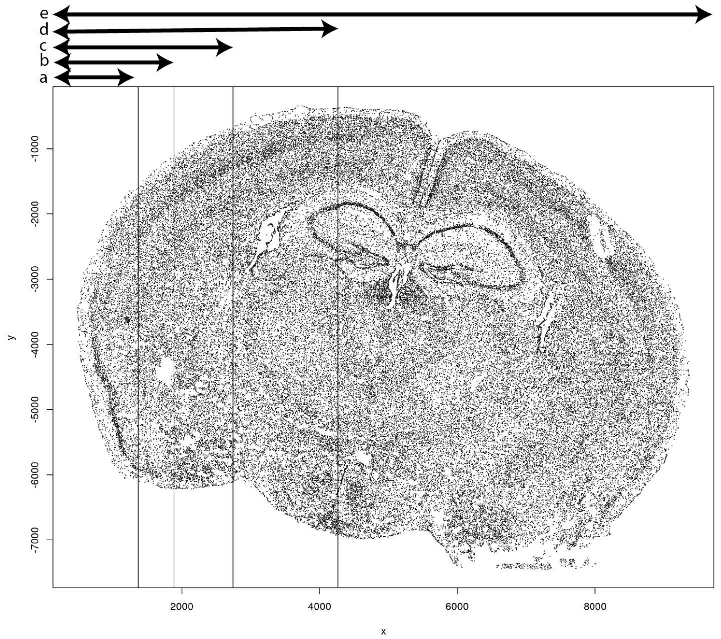


Figure 21: (i) Wall-clock time needed to run conventional clustering, BANKSY, MERINGUE and Giotto (HMRF) as a function of the number of cells. Plot vertical axis is in log scale. (ii) Tables showing the times; Top: overall run times for the four methods. Bottom three tables show the time taken for individual substeps of each method. (iii) Visualisation of the Vizgen dataset, slice 2, replicate 1. Vertical lines indicate the vertical strip used for each number of cells. For example, strip (c) extends from x-coordinate 0 to 2738, and takes all 16709 cells in this strip (which constitute 20% of all cells).
